# Emergent signal execution modes in biochemical reaction networks calibrated to experimental data

**DOI:** 10.1101/2021.01.26.428266

**Authors:** Oscar O. Ortega, Mustafa Ozen, Blake A. Wilson, James C. Pino, Michael W. Irvin, Geena V. Ildefonso, Shawn P. Garbett, Carlos F. Lopez

## Abstract

Mathematical models of biomolecular networks are commonly used to study cellular processes; however, their usefulness to explain and predict dynamic behaviors is often questioned due to the unclear relationship between parameter uncertainty and network dynamics. In this work, we introduce PyDyNo (Python Dynamic analysis of biochemical NetwOrks), a non-equilibrium reaction-flux based analysis to identify dominant reaction paths within a biochemical reaction network calibrated to experimental data. We first show, in a simplified apoptosis execution model, that Bayesian parameter optimization can yield thousands of parameter vectors with equally good fits to experimental data. Our analysis however enables us to identify the dynamic differences between these parameter sets and identify three dominant execution modes. We further demonstrate that parameter vectors from each execution mode exhibit varying sensitivity to perturbations. We then apply our methodology to JAK2/STAT5 network in colony-forming unit-erythroid (CFU-E) cells to identify its signal execution modes. Our analysis identifies a previously unrecognized mechanistic explanation for the survival responses of the CFU-E cell population that would have been impossible to deduce with traditional protein-concentration based analyses.

**Impact Statement:** Given the mechanistic models of network-driven cellular processes and the associated parameter uncertainty, we present a framework that can identify dominant reaction paths that could in turn lead to unique signal execution modes (i.e., dominant paths of flux propagation), providing a novel statistical and mechanistic insights to explain and predict signal processing and execution.

## INTRODUCTION

Many dynamic processes can be represented as networks of interconnected components such as the internet, ecology networks, social networks, and biochemical reaction networks. In cellular processes, networks comprising interactions between proteins, genes, and metabolites, have enabled the study of signaling and steady-state dynamics and their associated mechanisms (*1–5*). In systems biology applications, these studies typically entail building a graph that represents biochemical species (nodes) and biochemical interactions (edges), either from prior knowledge or through network inference, to then develop a mathematical representation for calibration and analysis of model parameters (*6–8*). Throughout the years, small networks with a handful of components have been studied with great success whereas the larger networks with several components and complex interactions present multiple challenges due to model parameter uncertainty. Depending on the network size, model complexity, i.e., the interdependency between the model parameters, and the amount of prior knowledge, some model parameters might be practically unidentifiable (*9*), which makes it very challenging to infer the underlying mechanism.

Multiple approaches have been used to extract mechanistic insights from mathematical models of network-driven processes. These approaches range from simple tracking of one or multiple species in the network (*10–14*) to more complex applications, such as Information Theoretic concepts to estimate the channel capacity between nodes in a network (*15–20*). Stoichiometric flux balance analysis methods are perhaps the most widely used approaches to analyze reaction flux in networks (*21–25*). These analyses allow computation of the steady-state flux distributions in biochemical networks by solving a constrained linear optimization (*26*) with the assumption that cells perform optimally with respect to a metabolic function of interest (*23*) such as growth or synthesis of a biomass (*27*), ATP production (*28*), and production of a specific metabolic product (*29*). Despite these advances to explore network-driven processes, they are typically applied to the systems at equilibrium. However, in real-world conditions, considering the dynamic interactions and flows of molecules, energy, and information within a network, equilibrium assumptions may not hold. Therefore, methods to analyze non-equilibrium network dynamics are of importance to gain a deeper understanding of emergent properties, system resilience, and the impact of perturbations. That said, a truly systems-level interpretation of non-equilibrium dynamics in a network and how these transient dynamics are impacted by parameter uncertainty remains a standing challenge in the field. Addressing this challenge might not only provide us with a better resolution of the signal processing mechanisms in complex biological systems but also enable us to construct biological hypotheses to be tested in the presence of parameter uncertainty.

In principle, the number of interactions and temporal dynamics present in a network makes it difficult to identify emergent behaviors from myriad concurrent biochemical reactions. We, therefore, developed PyDyNo, a Python-based dynamic network analysis tool, inspired in Tropical Geometry and Ultradiscretization Theory methodologies to map continuous functions into discrete spaces (*30,31*). The proposed dynamic analysis of biochemical networks enables us to identify dominant reaction fluxes that could in turn lead to emergent signal execution modes (i.e., unique dominant paths of flux propagation) and their dependence on model parameters. We cast this analysis onto a Bayesian probability framework to assign statistical weight to the identified modes of signal execution, thus providing a novel statistical and mechanistic interpretation for network-driven dynamic processes.

The remainder of the article is organized as follows. We first exemplify how model calibration can yield tens of thousands of parameter vectors that all reproduce an experimental data set equally well using Bayesian parameter inference on a simplified extrinsic apoptosis model. On this simplified model, we then introduce our method and define a dynamic signal execution fingerprint, which we then use to find whether execution modes can cluster despite the uncertain model parameters. Later, we demonstrate that choosing a single parameter vector offers a biased description of the signaling process, which could easily lead to misleading interpretations of execution mechanisms. Finally, we apply our approach to the Janus kinase 2/signal transducer and activator of transcription 5 (JAK2/STAT5) signal transduction pathway in erythroid progenitor cells, identify its signal execution modes that contribute to cellular survival responses upon stimulation with erythropoietin, and show how networks may switch their execution modes depending on the input dosage. Overall, our work introduces a probability-based methodology to explore non-equilibrium network-driven processes and to understand mechanistic model predictions in the presence of parameter uncertainty while providing an opportunity to gain novel insights into signal processing mechanisms in complex biological systems as elaborated in the subsequent sections.

## RESULTS

### Solutions from model optimization conflate protein concentration trajectories

To investigate the impact of parameter uncertainty on biochemical signal execution and gain mechanistic insights into non-equilibrium network dynamics, we present our method using a simplified early apoptosis execution model calibrated to experimental data using Bayesian parameter inference (*32*). Apoptosis is a ubiquitous biological process in metazoans used as a mechanism to maintain cell numbers and overall organism homeostasis (*33*). We explored the calibration of various versions of the Extrinsic Apoptosis Reaction Model (EARM) (*34*) to ensure that inferred model parameter values would converge to a distribution and could therefore be identified. The abridged EARM (abbreviated as aEARM in the rest of the paper), depicted in Figure 1A, was the largest model we could build that would both preserve key biochemical interactions and achieve convergence for all model parameters after model calibration. The abridged model preserves key biological features of apoptosis execution including signal initiation by TNF-Related Apoptosis Inducing Ligand (TRAIL), subsequent activation of initiator caspases (Caspase 8) (*35*), type 1 and type 2 activation of effector caspases (Caspase 3) (*36*), and completion of apoptosis execution by cleavage of Poly(ADP-ribose) polymerase (PARP) (*37*). Overall, aEARM comprises 22 molecular species and 34 kinetic parameters (see Materials and Methods for details).

**Figure 1.**
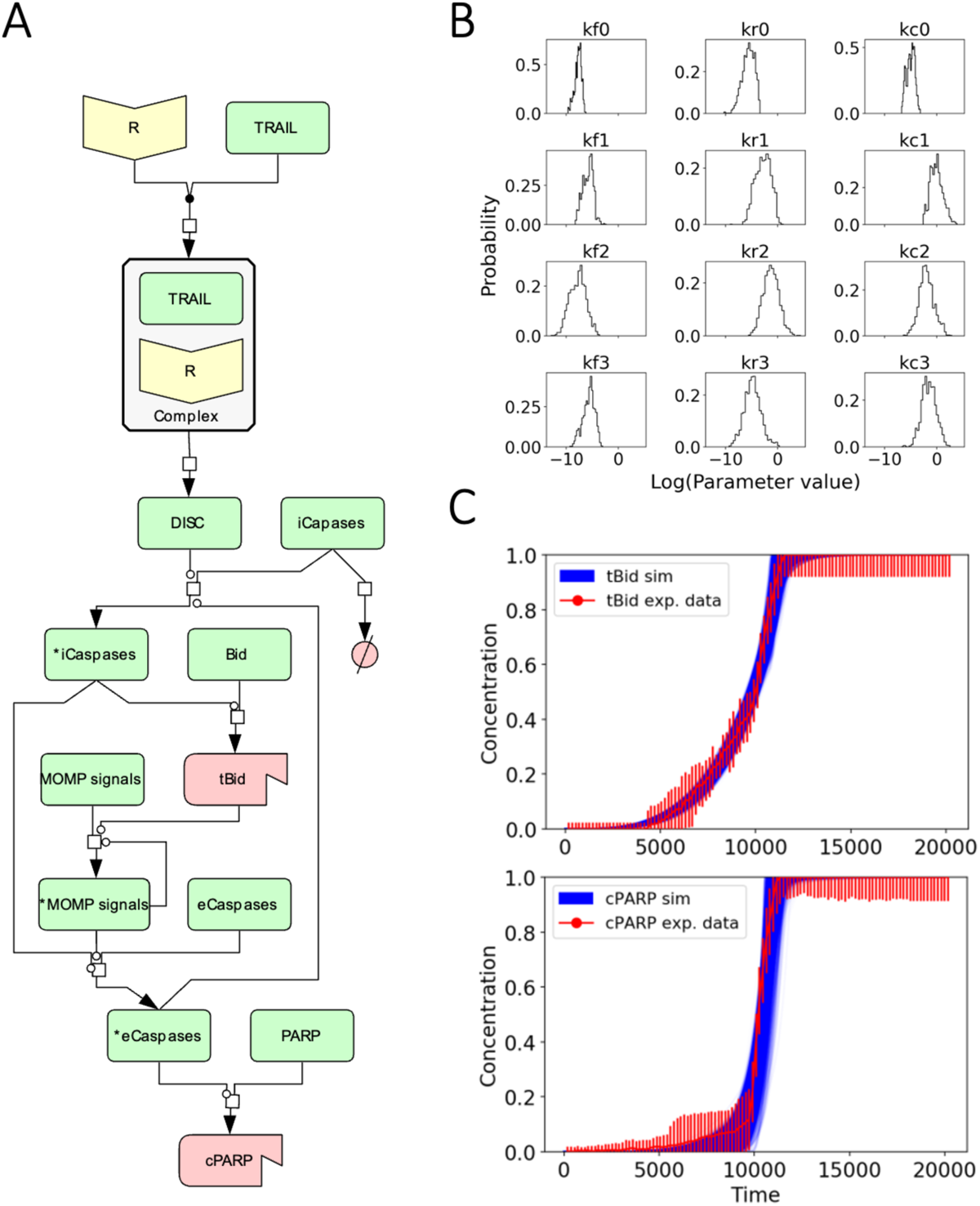
Abridged Extrinsic Apoptosis Reaction (aEARM) network and parameter calibration results. A) Reaction network using the Kitano convention. Yellow nodes are protein receptors, green nodes are generic proteins, and red nodes are truncated/cleaved proteins. B) Marginal probability distributions of the first 12 individual kinetic parameters that were recovered from the PyDREAM run by integrating all other dimensions. Forward rates, reverse rate, and catalytic values were all found to be within biologically relevant ranges (*39*). C) Simulated trajectories of truncated Bid and Cleaved PARP calibrated to reproduce the experimental data. Red dots and bars indicate the mean and standard deviation of the experimental data and blue lines correspond to the simulated trajectories.

We used PyDREAM – a python implementation of the MT-DREAM(ZS) algorithm (*8*) – to calibrate the model to previously published experimental data that comprises the concentration dynamics of truncated Bid (tBid) and cleaved PARP (cPARP) in HeLa cells (*38*). As HeLa cells are Type-II cells, the simulated results are representative of signal processing and execution of Type-II cells treated with death-inducing ligands such as TRAIL (*38*). The Bayesian model calibration yielded 27,242 unique parameter vectors that fit the experimental data equally well (see Materials and Methods). We note that throughout the manuscript, a *parameter vector* refers to a set of positive real values, one value for each of the kinetic parameters defined in aEARM, used to run a simulation. A parameter distribution refers to the frequency of occurrence of different values from the same kinetic parameter (Figure 1B, Figure S1) that provide equally good-fit to the experimental data (Figure 1C). Each model parameter distribution converged by the Gelman-Rubin diagnostics (*40*) (Table S1 and Figure S2). The probability distribution of parameter vectors exhibits characteristic exponential-like decay shape indicating that some parameters are more likely than others (Figure S3). Once we obtained a rigorously calibrated model, we then explored signal execution in aEARM from a probabilistic perspective.

### A discretized flux-based analysis of non-equilibrium signal execution in networks

As shown in Figure 1C, all parameter vectors obtained from Bayesian calibration yield protein concentration dynamics indistinguishable from the experimental trajectories of tBid and cPARP. Individual parameters from these vectors take widely different values as depicted by their distributions (Figure 1B). This uncertainty in the parameter values affects the reaction rates of the protein interactions generating different reaction flux patterns in the network during signal execution. We, therefore, studied the non-equilibrium flux of the reactions in the aEARM network to explore whether parameter uncertainty yields specific patterns of signal execution.

Analysis of flux dynamics during signal execution requires tracking the signal flow through a network at all simulation time points as multiple concurrent reaction rates consume or produce molecular species. For a particular species, we assume that the reactions with the highest flux at any given time dominate the network signal execution and provide a proxy to observe the effect of different parameter vectors in the network. Our aim is thus to identify the reaction rates with the highest flux throughout the whole network as simulations evolve. To analyze the non-equilibrium flux and find the dominant reaction paths during signal execution, we developed an algorithm inspired by Ultradiscretization Theory and Tropical Algebra (*31,41*) as described in the Materials and Methods section. This approach enables us to identify paths relevant for flux propagation in non-equilibrium states. We refer to these paths of flux propagation through the network as *execution modes* for the remainder of this manuscript.

We introduce the workflow for reaction flux discretization and execution mode identification as shown schematically in Figure 2. Signal discretization requires three steps. First, we identify a target node (Figure 2A) for which the signal flux will be tracked. Second, we calculate the reaction rates that produce or consume the target node, identify the largest reaction rate (*x*) and test whether it is dominant over other reactions (*y*) using the discretization operation |*x* | − |*y* | > ρ, where ρ is the order of magnitude difference necessary to consider dominance (see Materials and Methods). Third, we identify the chemical species produced by the dominant reaction(s) and jump to that species, thus starting the process again from the first step, and thereby tracking the dominant signal fluxes through the whole network and obtaining a subnetwork. This dominant subnetwork is assigned a unique integer label as shown in Figure 2A. The procedure is repeated for all simulation time points. As a result, the dynamic nature of signal execution for a given parameter vector is abstracted to a sequence of labels (Figure 2B) that can be compared to other sequences using a suitable metric. We call this sequence of labels obtained from a simulation as *dynamic fingerprint* because it is unique for a given signal processing event with a specific parameter set. The overall workflow of the algorithm is presented in Figure S4.

**Figure 2.**
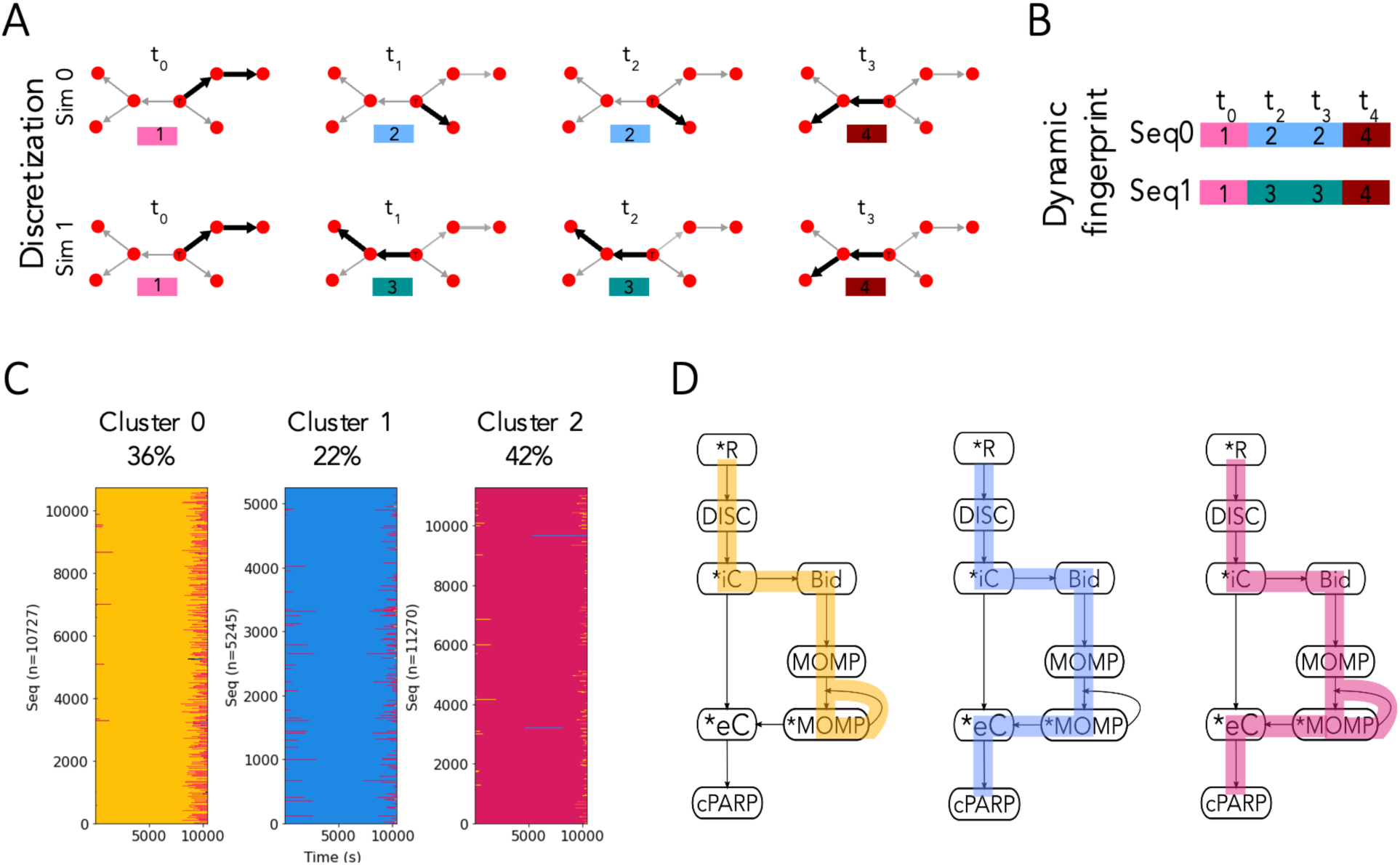
Discretization analysis workflow and modes of signal execution in aEARM. A) First, the network of interaction is obtained from a model and a target node (labeled T) from where the signal is going to be tracked is defined. Red nodes are molecular species in a model, edges represent interactions between nodes, and bolded edges are the dominant interactions. Next, at each time point of a simulation, our analysis obtains a dominant subnetwork, bolded edges in the network, through which most of the signal is flowing and this subnetwork is assigned a label. Sim 0 and Sim 1, simulations ran with different parameter sets, exhibit different dominant subnetworks. B) As each subnetwork is assigned a label, we can get a sequence of labels per simulation that can be compared to other simulations with the Longest Common Subsequence metric and obtain a distance matrix. This distance matrix can be used with clustering algorithms to obtain groups with similar modes of signal execution. C) Dynamic fingerprints organized by the clusters they belong to. Each cluster plot is composed of horizontal lines that correspond to dynamic fingerprints, i.e., sequences of dominant subnetworks, and each subnetwork is assigned a different color. D) Signal execution modes as defined by the most common subnetwork in each cluster. The complete aEARM network is shown in black, and the dominant subnetworks for Mode 1, 2, and 3 are highlighted in yellow, blue, and red, respectively.

### Key execution modes emerge despite parameter uncertainty

To identify the dynamic execution patterns in aEARM in response to death ligand cues, we carried out our signal discretization analysis for the 27,242 unique parameters and obtained dynamic fingerprints for each parameter vector. We then asked whether there were similarities among dynamic fingerprints across parameter sets. To investigate this question, we quantified the distance between each dynamic fingerprint using the Longest Common Subsequence (LCS) metric (see Materials and Methods) (*42*). We chose this metric due to its sensitivity to order differences in which successive subnetworks labels appear (*43*). This metric thus assigns a larger distance to a pair of dynamics fingerprints that execute the signal differently. Calculation of the pairwise distance between all dynamic fingerprints resulted in a 27,242 by 27,242 distance matrix. This matrix enabled us to use an agglomerative clustering algorithm (*44*) to probe whether clusters of dynamic fingerprints would emerge. As shown in Figure 2C, we found that all 27,242 dynamic fingerprints could be classified into three clusters (Table S2), which we denominate as “execution modes”. Given that each parameter vector has a defined probability (Figure S3) and is associated with a dynamic fingerprint, we calculated the probabilities of signal execution through each mode as 42%, 36%, and 22% for Execution Mode 1 (EM1), Execution Mode 2 (EM2), and Execution Mode 3 (EM3), respectively. These three execution modes account for all the parameter vectors inferred from the explored parameter space and no vectors were found that did not belong to either of these clusters. We note that these execution modes comprise three of the eight possible subnetworks for signal flow.

The dominant flux subnetwork for each execution mode is shown schematically in Figure 2D. The highlighted paths represent the dominant reaction fluxes, i.e., these fluxes are within an order of magnitude of the largest reaction at each node for the given parameter set and simulation time point. As shown in Figure 2D (yellow pathway), EM1 comprises events from initial death-ligand binding to the receptor, through the formation of the Death Inducing Signaling Complex (DISC), and subsequent activation of initiator Caspase (iC). The iC then truncates and activates Bid, which in turn activates MOMP, a species that abstracts mitochondrial outer membrane pore formation. As highlighted in Figure 2D (EM1), activated MOMP is dominantly used to both activate more MOMP, through the positive feedback loop, and activate the effector Caspase (eC). Note that although the species MOMP is represented as a single node in the model (Figure 1A), it indeed represents a more complex pathway through iCs, Bid, Bax, Smac, XIAP, and apoptosome which eventually leads to eC activation (*10*). Therefore, the direct interaction between MOMP and eC in the aEARM model is a simplification of the indirect interaction elaborated above.

The flux through the network in EM2 is similar to that of EM1 but the execution path differs at MOMP regulation. As highlighted in blue in Figure 2D (EM2), activated MOMP is largely consumed in the positive feedback loop to activate more MOMP. The signal flux downstream of activated MOMP is at least an order of magnitude less than the highlighted route for the parameters in EM2. Therefore, eC activation and apoptosis execution take place due to a smaller reaction flux in the network relative to the MOMP-level activity in EM2. For those parameters belonging to EM3, signal execution seems to flow largely toward PARP cleavage, with less MOMP-level regulation. Our results, therefore, show that despite uncertainties in inferred model parameters due to limited available data, the modes of signal execution are identifiable. Identifying a limited number of execution modes highlights the need to thoroughly characterize the model parameter space, given experimental constraints, to understand and make inferences about execution mechanisms. We note that using a single vector of parameters would lead to incomplete model prediction as no one single parameter vector captures the rich dynamics exhibited by all the statistically inferred parameter vectors.

Overall, the three execution modes arise from several parameter sets with equally well experimental data fitness. Although the difference between the dominant pathways in the observed execution modes might seem visually insignificant (i.e., a few interactions), they are indeed considerably different when considering the coarse-grained aspect of the aEARM model, as each execution mode is elaborated above. In addition, we would like to note that it does not mean that the nondominant pathways or interactions are not important or do not have any effect on the model responses, but rather it means that they are not the dominant interaction during the execution of the network process. For instance, eC is not included in the dominant pathway of EM2 (Figure 2D) while it exists in the other two modes. However, perturbing eC results in different responses in all execution modes compared to the wild type, which is thoroughly explained in the next section.

### Signal execution modes may respond differently to perturbations

We then asked whether *in silico* experiments could help us understand differences in signal execution that could lead to experimentally testable hypotheses. We, therefore, carried out *in silico* knockdown experiments of eC, as its activation is essential for the final steps of apoptosis execution (*45*). In addition, eC inhibitors are readily available for laboratory use (*46,47*). We tested the hypothesis that each execution mode would exhibit different execution mechanisms when eC is knocked down by 50%. To explore the impact of eC knockdown for each execution mode, we compared the concentration dynamics for MOMP and cPARP given by wild type (WT) and eC knockdown conditions.

For each execution mode, we plotted the cPARP concentration trajectories and obtained the time of death (ToD) for each simulated cell as described in Materials and Methods. As shown in Figure 3A, the ToD in EM1 exhibits a modest decrease of 14.96 s, but also presents a larger standard deviation of 702 s. For EM2, the ToD increases from 10351 ± 132 s (WT) to 10809 ± 226 s when eC is knocked down (Δt = 458 s). In contrast, EM3 eC knockdown leads to a decreased ToD from 10261 ± 83 in WT to 9507 ± 516 s in the knockdown (Δt = -754 ± 523 s). These results show that each execution mode can exhibit considerably different – and a time juxtaposed – responses to the same perturbation.

**Figure 3.**
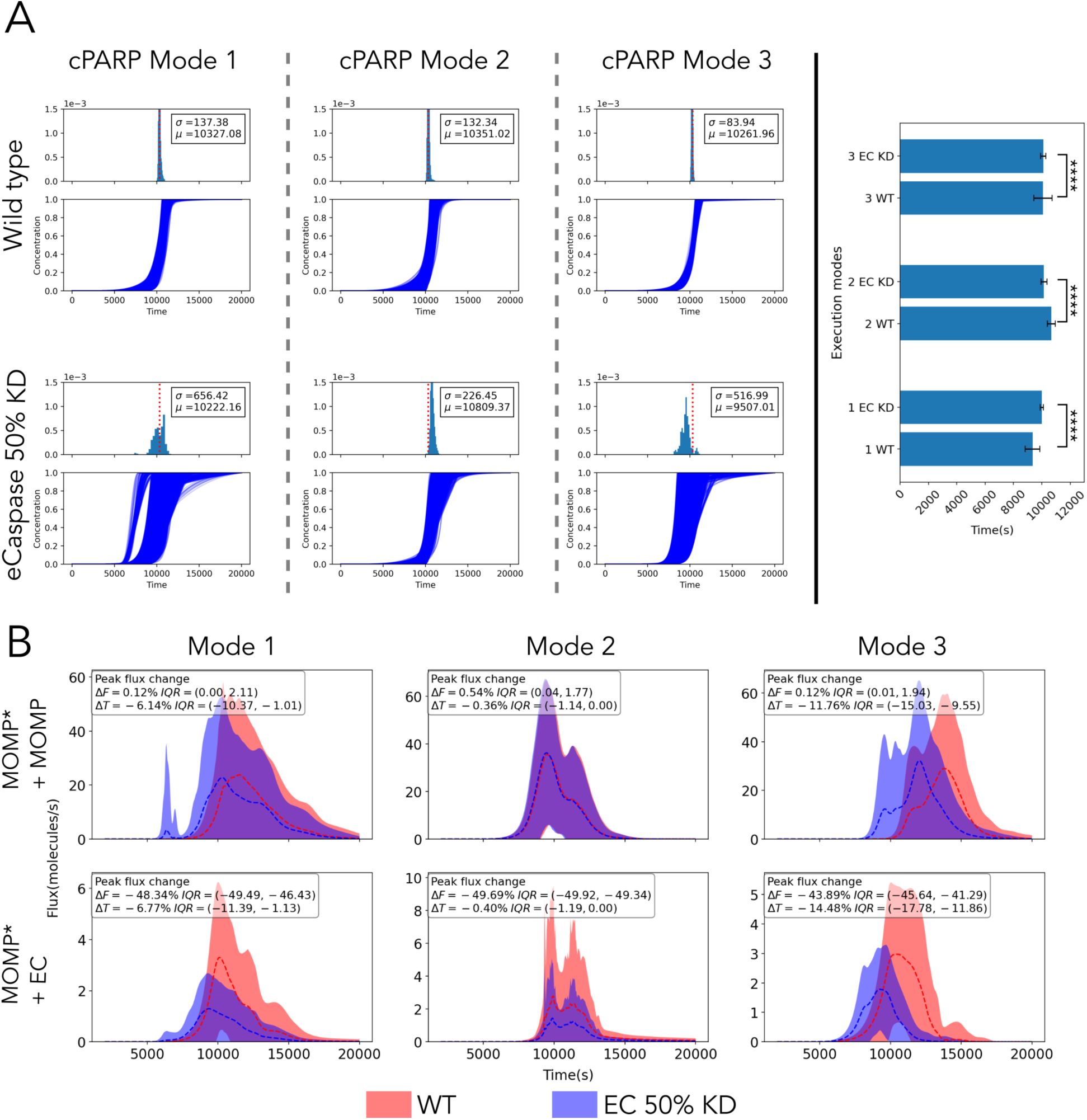
Time of death responses are markedly different for the same perturbation. A) Cleaved PARP (cPARP) protein concentration trajectories for the “wild type” case (top row) grouped by Execution Modes. Mode 1, 2, and 3 have 11270, 10727, and 5245 trajectories, respectively. Inset includes the average time to death and the standard deviation calculated from all trajectories in each execution mode. PARP cleavage exhibits a markedly different trajectory pattern (bottom row) after eCaspase is knocked down by 50%. B) MOMP* + MOMP and MOMP* + eC reaction rate trajectories. Dashed lines correspond to the mean of all reaction rates trajectories in an execution mode and the shadows represent the standard deviations. Trajectories from the “wild type” condition are colored in red and trajectories from the 50% effector caspase KD are colored in blue, and show key differences in their dynamics. Insets include the median percentage change in the reaction rate peak (ΔF) and the time to reach that peak (ΔT) in the eC KD condition relative to the wild-type condition. The interquartile range is included as a measure of the variation in the ΔF and ΔT changes.

We then probed the effect of the eC knockdown on the reaction rates associated with MOMP* (a node where the signal bifurcates): MOMP* binding to MOMP, and MOMP* binding to eC. Specifically, we focused on the reaction rate peak and the time to reach peak of the reaction rate throughout the simulation, as shown in Figure 3B. The peaks of the MOMP*+MOMP binding reaction (Figure 3B upper row) appear unchanged across all execution modes, yet the time to reach the peaks varies significantly. Compared to the WT simulations, the median time to peaks is 6.14%, 0.36%, and 11.76% faster for modes 1, 2, and 3, respectively. Concurrently, the peaks of the MOMP*+eC binding reaction (Figure 3B lower row) reduce approximately 50% as expected by the 50% reduction of the available eC, and the median time to peaks are 6.77%, 0.4%, and 14.48% faster for modes 1, 2, and 3, respectively. In combination, for mode 1, the relative change of the MOMP and eC reaction peaks have large interquartile ranges IQR= – 10.37% to –1.01% and IQR=-1.13% to –11.39%, respectively, which explains the variability in the time to cell death. For mode 2, the time to the peak of MOMP and eC reactions change marginally and given that the eC peak is 50% of the WT condition, this leads to longer times to accumulate the necessary number of eC molecules for cells to commit to apoptosis. Finally, for mode 3, the median time to reach the MOMP reaction peak and the eC reaction peak are 11.76% and 14.48% faster than in the WT condition, respectively. This causes faster activation of MOMP and eC which leads to earlier apoptosis in cells.

In addition to perturbation analysis of the execution modes, we showed how the execution mode uncertainty can be reduced through parameter measurements using machine learning techniques in the Supplementary Materials (Section 1). Furthermore, we studied the signal execution modes of the extended apoptosis model (EARMv2.0) (*34*) to show the feasibility of the proposed framework on a larger model with higher parameter uncertainty as well as how the network size affects the identified execution modes, whose details are also provided in the Supplementary Materials (Section 2).

To summarize, although the biochemical signal flows differently in each execution mode, the protein concentration dynamics exhibit similar outcomes (Figure 3A Wild Type). However, when a perturbation is made to the network, the outcome can vary significantly, as shown for each execution mode. Considering that each execution mode was obtained from several parameter vectors with equally well data fitness, such a difference is theoretically noteworthy and might be an indicator of a more significant difference in other systems. Moreover, these observations highlight the importance of choices of model parameter values as different parameter vectors may result in different outcomes. Therefore, care should be taken while analyzing data-driven network models in the presence of parameter uncertainty.

### Networks may switch their execution modes depending on the input dosage: Application on JAK2/STAT5 signal transduction network

To show how the proposed dynamic network analysis pipeline can provide mechanistic insights into other biological systems, we applied it to a model of the Janus kinase 2/signal transducer and activator of transcription 5 (JAK2/STAT5) signal transduction pathway that captures cellular population response (*48,49*). This pathway has an active role in the survival of erythroid progenitor cells (*50*), which are highly sensitive to erythropoietin (Epo) at the colony forming unit erythroid (CFU-E) stage (*51*). To ensure a robust physiological production of erythrocytes, these cells demonstrate a gradual input-output relationship when exposed to Epo (*52*). The details of the JAK2/STAT5 signaling network population-average model and its calibration, the data used in the model calibration, and the uncertain parameter vectors providing equally well data fitness were obtained from Adlung et al. (*49*).

The JAK2/STAT5 signaling network is presented in Figure 4A. In summary, the signal transduction, which eventually induces gene expression (*53*) and CFU-E survival (*48*), is initiated by the binding of Epo to its receptor EpoR. This is followed by the activation of receptor-associated JAK2 by the formed Epo-EpoR complex. Active JAK2 phosphorylates both the cytoplasmic tail of EpoR on multiple tyrosine residues (*54*) and STAT5 that is recruited into the receptor complex (*55*). Then, phosphorylated STAT5 (pSTAT5) translocates to the nucleus and induces the anti-apoptotic genes for the control of cell survival (*56*). STAT5 also targets other genes including cytokine-inducible SH2-domain-containing protein (Cish) (*57*) and suppressor of cytokine signaling 3 (SOCSC3) (*58*), which attenuates signal transduction. Finally, signal transduction is terminated by dephosphorylated JAK2 (*54*) caused by the recruitment of SH2-domain-containing protein tyrosine phosphatase 1 (SHP1) to the activated receptor via its SH2 domain (*59*).

**Figure 4.**
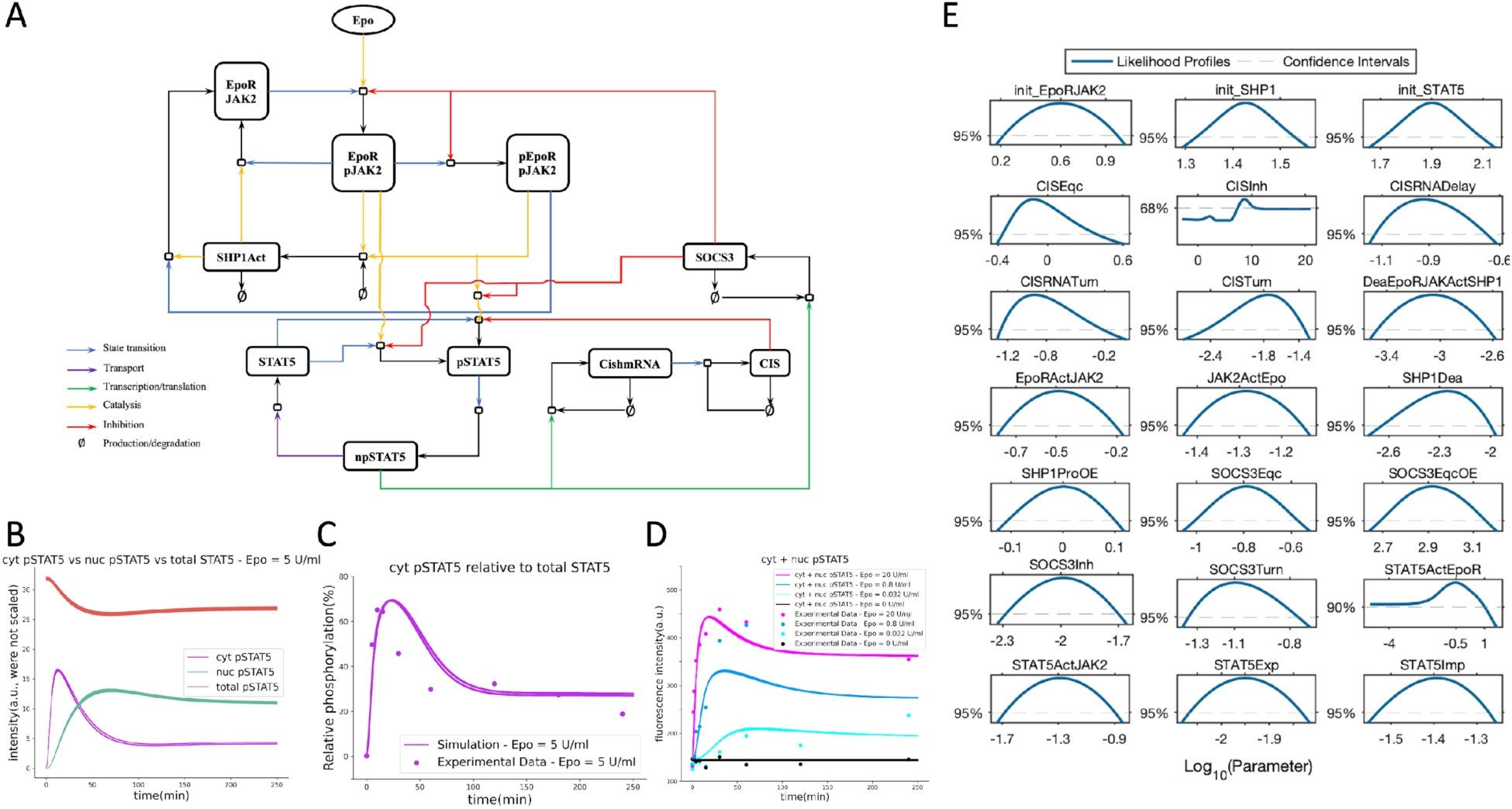
JAK2/STAT5 population-average model, model parameters, and STAT5 responses reproduced using the calibrated model provided by Adlung et al. (*49*). A) Graphical representation of the JAK2/STAT5 signaling network model. B) Comparison of average total STAT5, pSTAT5 in the cytoplasm (cyt), and pSTAT5 in the nucleus (nuc) responses of CFU-E cell population over time when exposed to Epo = 5 U/ml. C) Simulated pSTAT5 (cyt) relative to total STAT5 versus the experimental measurements. D) Overall pSTAT5 responses over time with respect to increasing Epo doses versus corresponding experimental data. Note: The experimental data and the model parameters were provided by Adlung et al. (*49*). 1907 parameter sets were simulated in total and plotted all together. As seen in B) and C), all parameter sets are fitting the data equally well and their trajectories are mostly overlapping. E) Likelihood profiles of the 21 dynamic model parameters (*49*).

Although the fraction of surviving cells increases at a graded level, the population mean fluorescence intensity of the total STAT5 – the key integrator of Epo-induced survival signal transduction in CFU-E cells – does not change with respect to increasing Epo doses (*48,49*). Upon exposing the cells to Epo = 5 U/ml, the total STAT5 remains nearly constant (Figure 4B). On the other hand, the mean fluorescence intensity of the pSTAT5 in the population gradually increases in cytoplasm and nucleus over time which correlates with survival responses after Epo stimulation (Figure 4B). Similarly, CFU-E cells demonstrate a rapid increase in the level of pSTAT5 in cytoplasm relative to total STAT5 (Figure 4C). Furthermore, the overall pSTAT5 levels increase as the Epo level increases (Figure 4D), which is correlated with the increasing cell survival (*49*).

Despite the model reproducing the experimental data and identifying the abovementioned key regulators of CFU-E cell survival responses with respect to Epo doses (*49*), a mechanistic understanding of these networked processes that give rise to this behavior was lacking. Revealing the underlying mechanism of this behavior and understanding of signal processing mechanism of CFU-E cells in the presence of parameter uncertainty would enable us to control their population-level responses through their signal execution modes. This could pave the way for model-driven hypothesis generation to control cellular states and state transitions. Therefore, here, we employ this system with the provided parameter distributions (Figure 4E) and apply our proposed dynamic network analysis framework to unveil the underlying mechanism of this behavior by identifying the JAK2/STAT5 network execution modes. Moreover, we analyze how the number of modes varies with respect to the increasing Epo levels and demonstrate the time-dependent dominant pathways that play a role in the increase of pSTAT5 levels (Figure 5B) and hence the cell survival responses.

**Figure 5.**
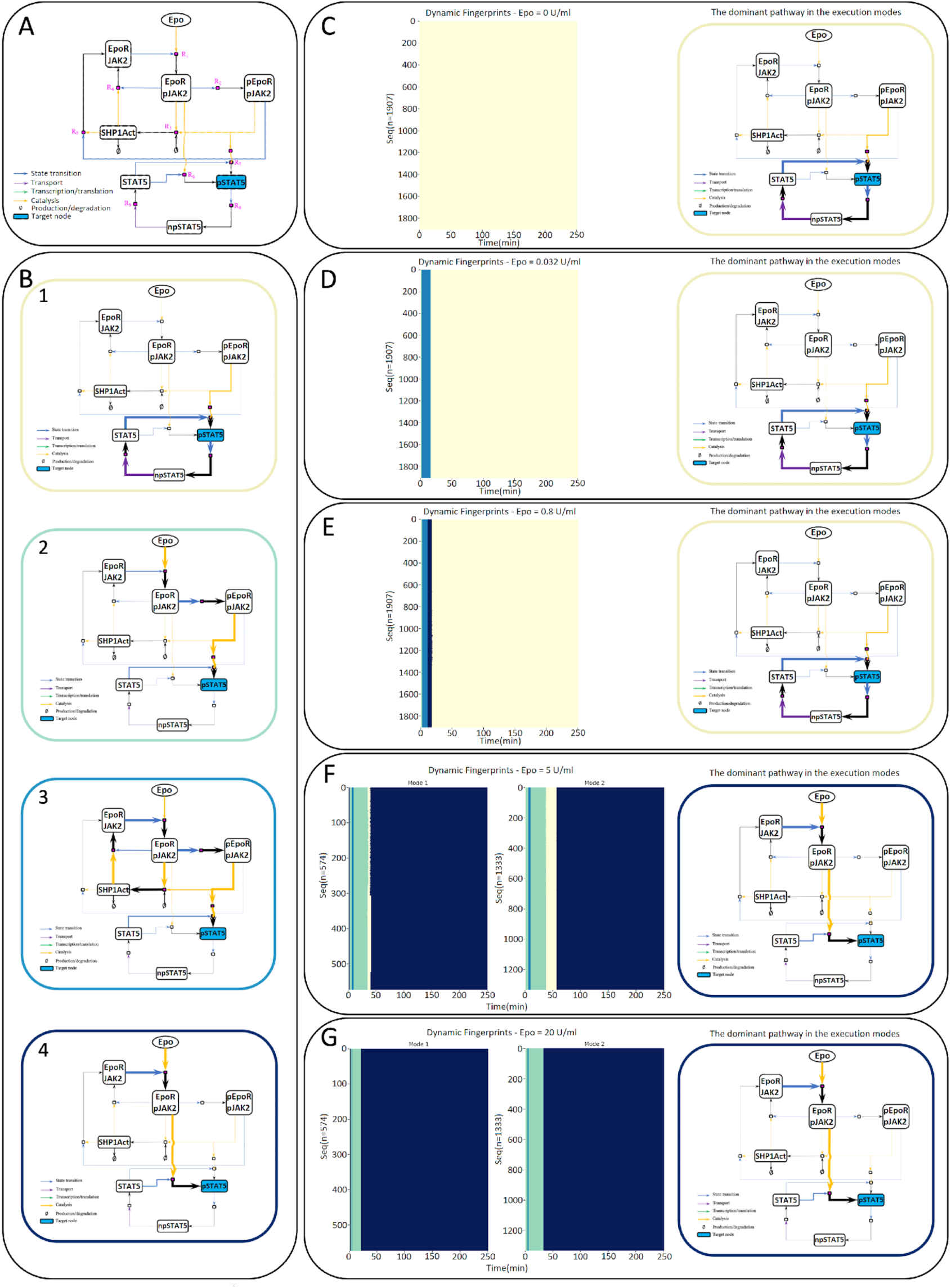
Execution modes of JAK2/STAT5 signaling network for each Epo dose when pSTAT5 is the target node. A) The pruned JAK2/STAT5 signaling network. The inhibitory interactions were removed for demonstration purposes to show the positive reactions only. B) Dominant pathways over time framed with the corresponding colors in the identified execution modes in C-G. C-G) Dynamic fingerprints of the network for each parameter set (each row corresponds to a fingerprint sequence for a unique parameter set) when Epo = 0 U/ml, 0.032 U/ml, 0.8 U/ml, 5 U/ml, and 20 U/ml with their most common dominant pathways, respectively.

### The JAK2/STAT5 signaling network execution modes and the dominant pathways over time vary after stimulating with different Epo doses, which correlate with the cell survival rates

To apply the proposed dynamic network analysis framework to the JAK2/STAT5 signaling network, we first implemented the model in Python using the PySB modeling tool (*34*), which was further used to reproduce the network responses (Figure 4B-D). The model consists of 21 dynamic parameters as well as several offset and scaling parameters, in which the parameters have uncertainty with wide likelihood-based confidence intervals (Figure 4E) and equally good fitness to experimental data (*49*) (Figure 4C-D). Among the provided parameter distributions, we picked the best-fitting 1907 parameter vectors and used them to identify the execution modes.

First, we identified pSTAT5 as the target node because we are interested in the mechanism that causes the increase of pSTAT5 in both cytoplasm and nucleus which is correlated with the cell survival responses. For illustration purposes, we pruned the JAK2/STAT5 signaling network by removing inhibitory interactions and created the production network highlighting only positive reactions (Figure 5A). Next, we computed the reaction rates that produce pSTAT5 and tracked the signal flux to create the dominant subnetwork at each time point. Then, we labeled the identified unique dominant subnetworks by a unique integer and obtained the dynamic fingerprint of the network. Lastly, we repeated these steps for each parameter vector. After computing similarities between each obtained sequence using the LCS metric and clustering them, we observed the different signal execution modes of the network for each Epo dose (see Materials and Methods section for more details on each step).

Interestingly, JAK2/STAT5 network switches its execution mode based on the Epo dosage (Figure 5C-G), in which various pathways become dominant over time (Figure 5B). When Epo doses are low (e.g., Epo = 0, 0.032, and 0.8 U/ml), the signal is executed in a single mode, but the modes are slightly different (Figure 5C-E). More precisely, when Epo = 0 U/ml (Figure 5C), the only dominant pathway that can induce pSTAT5 production over time is the feedback loop between pSTAT5, npSTAT5, and STAT5 (Figure 5B-1), which depends on the initial STAT5 abundance only. When Epo = 0.032 U/ml (Figure 5D), three different pathways that become dominant at different time intervals emerge. To elaborate, at t = 1 min after Epo was applied, JAK2 is activated by the Epo-EpoR complex, and eventually phosphorylated receptor complexes (i.e., EpoRpJAK2 and pEpoRpJAK2) are formed. Then, upon the involvement of STAT5 with pEpoRpJAK2, initial levels of pSTAT5 are observed (Figure 5B-2). Between t = 2 – 15 min, a cytosolic protein, SHP1, is recruited into the active receptor complexes and dephosphorylates JAK2 to terminate signal transduction (Figure 5B-3). Between t = 16 and 250 mins, the signal from medium to cytoplasm and nucleus attenuates and the only pSTAT5 production source becomes the STAT5 feedback pathway (Figure 5B-1). Similarly, when the Epo dose is increased to 0.8 U/ml from 0.032 U/ml, the network executes the signal in a single mode with slightly different dominant pathways (Figure 5E). This increase in Epo level reduces the duration of the activity of signal transduction terminator pathway from 14 to 9 mins and additionally activates one of the direct pSTAT5 producer pathways (Epo–EpoRpJAK2–pSTAT5; Figure 5B-4) which contributes to the production of more pSTAT5 for 7 mins. However, this activation does not last long due to weak Epo dose and again the STAT5 feedback loop becomes the only dominant pathway producing pSTAT5 between t = 18 and 250 min.

Increasing the Epo level not only substantially changes the network’s execution mode but also further increases the number of modes and changes both order and duration of the dominant pathways (Figure 5F-G). When Epo = 5 U/ml, the dominant signal flux begins with the activation of the direct pSTAT5-producing pathway (Figure 5B-4) at t = 1 min and switches to the other pSTAT5-producing path through the phosphorylated receptor complex pEpoRpJAK2 (Figure 5B-2) between t = 2 and t = 34 min in the first mode. In the meantime, although the signal transduction terminator pathway through SHP1 becomes dominant for 4 mins, it is not enough to prevent pSTAT5-producing pathways from becoming dominant again because of the strong Epo signaling. First, the STAT5 feedback pathway becomes dominant between t = 35 and t = 39 min and then it is followed by the major dominance of the direct pSTAT5-producing pathway (Figure 5B-4) from t = 40 mins until the end of the simulation, maintaining high levels of pSTAT5. The same scenario occurs in the second execution mode as well. The only difference between the two modes is the duration of the dominance of the STAT5 feedback pathway (Figure 5F, Mode 2). When Epo is applied at the level of 20 U/ml (Figure 5G), both execution modes have the same dominant pathways that directly produce cytoplasmic pSTAT5 (Figure 5B-2 and 4). Only the duration of the dominance of the pathway through the pEPoRpJAK2 receptor complex lasts longer in the second mode, which results in barely observable, long-lasting pSTAT5 peak levels.

The change in the signal execution modes and the temporal dominant pathways with respect to increasing Epo doses is consistent with the increase in the cell survival percentages. When the Epo level ranges from 0 to 20 U/ml, the percentage of surviving cells gradually increases from 6.9% to 82%, inferring a correlation with the high responses of pSTAT5 (Figure 4D) (*49*). As elaborated above, increasing the Epo level results in quantitatively and qualitatively different signal execution modes. Comparing all signal execution modes in Figure 5C-G indicates that when Epo levels are low, the main dominant pathway contributing to the pSTAT5 production is the STAT5 feedback pathway in which the production rate mostly depends on the initial abundance of STAT5 and is weakly affected by the Epo-induced signaling, resulting in lower cell survival rates (Figure 5C-E). On the other hand, when the applied Epo dose becomes higher, the cells mechanistically switch to another state. More precisely, the main dominant pathways over time become the direct pSTAT5-producing pathways through Epo receptor complexes and phosphorylated JAK2 (Figure 5F-G), which results in a more robust pSTAT5 production and hence higher cell survival rates. The relationship between varying Epo doses, the signal execution modes, and the resulting pSTAT5 trajectories are further demonstrated in Figure S5.

As a result, despite the parameter uncertainty, by applying the proposed framework to the JAK2/STAT5 signal transduction pathway model and identifying its switch-like signal execution modes, we not only reveal novel, detailed explanations of how different levels of Epo signaling are mechanistically processed by the CFU-E cells but also show how the parameter changes, i.e., the change in applied ligand dosage, etc., may affect the flux and signal execution in networks. We hypothesize that these Epo-dependent execution modes might be further used to precisely control CFU-E cell population level decisions by specifically targeting the associated dominant pathways through experiments. Furthermore, this analysis shows that the proposed approach might be used to reveal broad information about signal flow in the other complex systems and perhaps to elicit ways of reprogramming the cells and controlling their fates through their unique signal execution modes.

## DISCUSSION

It has been long recognized that model parameter optimization to experimental data is key to investigating the dynamical properties that control cell behavior (*60*). Unfortunately, parameter optimization usually yields large parameter uncertainties due to a general lack of quantitative data, model complexity as well as model identifiability (*61*). Even a complete set of time course data might be insufficient to constrain parameter spaces, yielding unidentified parameters (*62*). The unidentifiability of model parameters refers to a situation where it is not possible to uniquely estimate the values of individual parameters based on the available data. This can occur when different combinations of parameter values produce similar model predictions or when certain parameters have no direct influence on the model output. However, while individual parameters may be unidentifiable, combinations of model parameters can often exhibit identifiability in which identifiability means the ability to estimate the values of specific combinations of parameters with a high degree of confidence. By examining the relationships between parameters and their influence on the model output, it is possible to identify specific combinations that have a discernible impact on the system behavior. These combinations may represent key regulatory mechanisms, interaction networks, or functional modules within the model.

Given the set of parameter vectors yielding equally well data prediction accuracy, analysis of non-equilibrium network dynamics is of importance because it allows us to capture and study the behavior of complex systems in real-world conditions, where equilibrium assumptions may not hold. By considering the dynamic interactions and flows of molecules, energy, and information within a network, one can understand emergent properties, system resilience, and the impact of perturbations. This knowledge is not only essential for biological questions, but also essential in various fields such as physics, ecology, and social sciences, as it provides insights into complex processes, enabling us to make informed decisions and develop effective strategies in complex, non-equilibrium systems. Here, we examined the effects of parameter uncertainty on signal execution through a biochemical network with an approach inspired by Tropical Geometry and Ultradiscretization Theory, called PyDyNo. Despite the many parameter vectors which reproduce the experimental protein dynamics, we found that the signal flow in a network was constrained to only a few modes of execution. Our analysis further shows that within a Bayesian calibration scheme, it is possible to assign probabilities to each execution mode, thus greatly improving our understanding of signal dynamics. Therefore, the probabilistic approach introduced in this work could open a novel perspective to understand network-driven processes from a statistical point of view.

To introduce the proposed framework, we first studied a simplified extrinsic apoptosis reaction model (aEARM) that comprises key biological properties of apoptosis and allows us to analyze all aspects of our method thanks to the convergence of all model parameters after parameter calibration. Despite the found 27,242 unique parameter vectors with equally good data fitness, only three distinct execution modes emerged as signal processing mechanisms, which exhibited considerably different responses to *in silico* perturbations, indicating that using a single best-fit parameter vector could be insufficient and misleading for understanding signal dynamics in complex models. In addition to the aEARM, we studied the signal execution modes of the extended apoptosis model (EARMv2.0 in Supplementary Materials) that provides a more detailed resolution to the apoptosis mechanism, to see how the network size affects the execution modes and overall prediction. Despite the difference in the underlying system of differential equations due to the resolution of the models, the identified highly probable execution modes share a functional overlap between the underlying dominant pathways, which may imply that the execution modes are intrinsic features of the underlying system and independent of the specific way in which the model is implemented. In addition, we applied the framework to JAK2/STAT5 signal transduction network in erythroid progenitor cells to reveal a mechanistic explanation for their survival responses with respect to increasing erythropoietin (Epo). Upon applying different Epo doses, we observed that the network switches from one execution mode to another execution mode resulting in weak to robust cell survival rates.

Although our approach provided novel insights about signal execution in important biological networks, it has certain limitations. Our analysis assumes that reactions with high flux are the most important for signal processing in a network. However, this may not always be the case for other networks or for networks with temporal changes in model topologies (*63*). Moreover, our method could be computationally expensive, particularly as models increase in size, requiring hundreds of thousands of parameter samples to reach a convergence criterion. Nevertheless, we believe this is a relatively small price to pay in contrast to the number of experiments that would be necessary to attain the same level of mechanistic knowledge about a network-driven process.

Overall, the proposed framework is advantageous in the non-equilibrium analysis of data-driven complex network models by providing broad information about signal flow and how the system of interest mechanistically executes the signals in the presence of parameter uncertainty. In addition, the knowledge of the dominant pathways contributing to the production (or consumption) of a target protein over time could guide the researchers through experiments and allow easy identification of the other experimental targets to test hypotheses about the target protein and the pathway itself. Also, the identification of signal execution modes might guide how to precisely control the cell behaviors, cellular states, state transitions, and perhaps their fates when they are exposed to an input signal. Furthermore, the information about signal flow could be used to study drug-induced network rewiring processes (*64*), provide mechanistic explanations for drug responses, and predict sequential combinations of drugs that could better modulate a response signal in biochemical networks.

## MATERIALS AND METHODS

PyDyNo (Python Dynamic analysis of biochemical NetwOrks) is a dynamic network analysis tool, inspired by Tropical Geometry and Ultradiscretization Theory methodologies to map continuous functions into discrete spaces. In short, the algorithm works as follows: given a mechanistic model of a network-driven process with associated parameter sets that yield equally well data prediction, for each parameter set, we first discretize the simulated trajectories at each time point and identify the dominant signal fluxes through the whole network and observe a subnetwork. Upon labeling each subnetwork at all the time points, we abstract the dynamic nature of signal execution to a sequence of labels, which is called as dynamic fingerprint. Clustering of the dynamic fingerprints obtained from each parameter set results in the execution modes. Each of these steps is elaborated in the following subsections and the overall workflow of the algorithm is presented in Figure S4.

### 1. Development of the abridged extrinsic apoptosis reactions model (aEARM)

A coarse-grained ODE model of TRAIL-dependent apoptotic signaling was encoded with PySB (*34*). This model grouped reactions into simple dynamic motifs representing key mechanistic blocks in the TRAIL-dependent apoptosis pathway: TRAIL mediated DISC formation, initiator caspase-8 activation via the DISC, and feedback activation via effector caspases (-3, -6, and -9), effector caspase activation, and apoptotic marker (PARP) cleavage were all encoded as simple catalysis reactions. MOMP formation (via initiator caspase-activated Bid) and accumulation of MOMP-dependent pro-apoptotic signals were modeled as a Bid-dependent activation and subsequent feedback self-activation step. This activation-amplification motif reproduces the observed sigmoidal “snap-action” dynamics of MOMP-dependent pro-apoptotic effectors (e.g., Smac, CytoC). Initial values of the model components were drawn from values present in earlier apoptosis models (e.g., EARM). Initial values of the MOMP-dependent signaling component took the same value as Bax and Bak in the EARM model. The ODE model was integrated using the Python LSODA ODE solver with relative and absolute tolerances set to 1e-2 and 1e-1 respectively (the model is encoded in copies per cell which takes values of 10,000 to 1M).

### 2. Bayesian inference and parameter calibration

The model’s reaction rate parameters were calibrated to normalized iC-RP and eC-RP fluorescence time-course data, using the Differential Evolutions Adaptive Metropolis MCMC sampling algorithm (DREAM(ZS)) encoded in Python as PyDREAM. This Bayesian calibration uses log-normal distributions as priors, centered at biologically plausible rate values of forward binding, reverse binding, and catalysis (1e-6 s^-1^ molecule^-1^, 1e-3 s^-1^, 1 s^-1^). The likelihood function assumes the iC-RP and eC-RP data are normally distributed with a mean and standard deviation calculated from multiple measurements. The sampling process employed a burn-in of 80,000 steps followed by a 220,000-step sampling of the target distribution. Additional settings were applied to the gamma term in the DREAM algorithm: number of crossovers (nCR) = 25, adapt gamma = True, probability of gamma-unity (p_gamma_unity) = 0.1, resolution of gamma term (gamma_levels) = 8. Convergence was diagnosed by the Gelman-Rubin convergence diagnostic (i.e., GR ≤ 1.2) for each calibrated parameter. This calibration provided a wide range of kinetic parameter values, and we note that even if there was experimental data for all species in a model, due to model sloppiness the parameter distributions would not be sufficiently constrained (*62,65*).

#### a. Calculating time of death from simulations

To calculate the time of death from a simulation, we used the same method used by Lopez et al. (*34*). This calculation consists in obtaining the times taken by the simulated trajectory of cPARP to reach 10% and 90% of the maximum concentration. Then the time to death is calculated as follows:

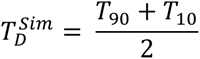

### 3. Constructing the digital signature for a subnetwork

The digital signature of a model simulation is a temporally ordered sequence of labels denoting the dominant sub-network at each time point of the simulation. The dominant sub-network for a given time point is a subset of the signaling pathway over which the most signal is flowing to the production of a pre-determined target species (i.e., signaling protein or protein complex); each unique dominant sub-network is assigned a unique integer label for the digital signature.

Construction of the dominant sub-networks uses the instantaneous reaction rates at each simulation time point, which change over time as molecular species are consumed and produced. Hence, the dominant sub-networks can change at each time of the model simulations, resulting in a dynamic sequence of dominant sub-networks, i.e., the digital signature of the model simulation. The procedures for constructing the dominant sub-networks and for selecting the dominant reaction are detailed in the proceeding paragraphs. Figure S4 shows an algorithm diagram for the construction of digital signatures.

#### a. Constructing a dominant sub-network

The generation of the dominant sub-network at a given time point starts with creating a bipartite-directed graph (digraph) representation of the model which consists of molecular species and reaction nodes. All molecular species and reactions within the model are encoded as nodes with unidirectional edges connecting molecular species with their respective reaction nodes. The directionality of each edge is determined by the sign of the reaction rate of the given species-reaction pair at the current time point; for reversible reactions, the reaction rate is the sum of forward and reverse rate terms. The bipartite digraph provides an instantaneous snapshot of the direction of fluxes through the signaling network.

We then hierarchically construct a dominant pathway starting from the user-defined target species. For the first step, we identify the set of dominant reactions, i.e., those reactions which contribute most to the production of the target species. Dominant reactions are classified based on the instantaneous reaction rates; the conditions determining the dominant reactions are detailed in the following paragraph. We then trace back through the bipartite graph along those dominant reactions to the corresponding reactant species, which are added to the dominant sub-network. For each reactant species that was added to the sub-network, we determine their dominant reactions and trace back through the bipartite graph to the next set of reactant species. This procedure is continued for a pre-determined number of iterations defined by the user parameter *depth*. Obviously, the length of the identified dominant pathway at each time point depends on the user-defined depth value. Using higher depth values would result in longer dominant pathways with more species while making the algorithm computationally more complex. Once the procedure is complete, the result is a species-to-species sub-network representing the dominant pathway over which most of the signal is flowing to produce the target at the current time point.

Note that we have defined the dominant sub-networks and their construction based on *production* of the target species. Alternatively, the procedure can be formulated to define the dominant sub-networks and their construction based on the *consumption* of the target species.

#### b. Selecting dominant reactions

One aspect of network complexity is that signaling proteins can participate in multiple interactions; in a bipartite-directed graph, this is represented by a molecular species having edges connecting it to multiple reactions. The goal, therefore, is to simplify the system and focus on only those reactions which are most important to the production of a species; we term this sub-set of reactions the dominant reactions. To determine the set of dominant reactions we build on some of the concepts from Noel et al. (*31*), who applied a tropical geometry framework to smooth ODE systems.

When the signaling network is modeled with a system of ODEs the rate of change of a molecular species, *x_i_*, is:

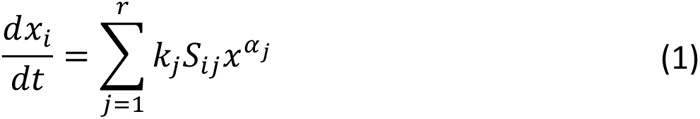

where *k_j_* > 0 are kinetic constants, *x_i_* are variable concentrations, 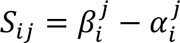 are the entries of the stoichiometric matrix, 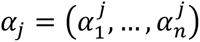 are multi-indices, and 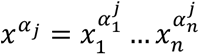. We treat the ODE of molecular species x*_i_* as a polynomial function of the uni-directional rate terms,

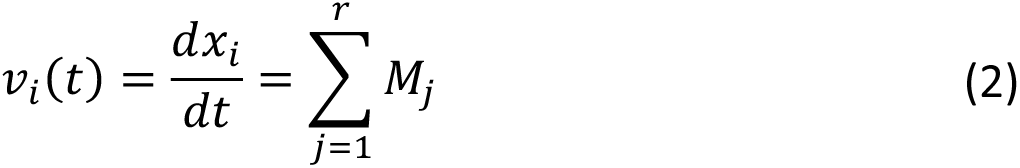

where the M_j_ = *k_j_S_ij_x^aj^* are the monomials representing the uni-directional rate terms. Since reversible reactions are bi-directional, they contribute two uni-directional rate monomials to the total rate of change of a species: one each for the forward (consumes *x_i_*) and reverse (produces *x_i_*) directions of the reaction. We account for this bi-directionality by combining the two uni-directional rate monomials into a single net reaction rate monomial term,

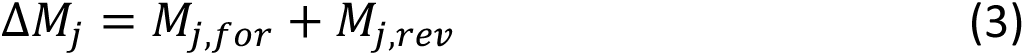

where *M_j,for_* and *M_j,rev_* are the monomials corresponding to the forward and reverse reaction rate monomials of reaction *j* with respect to species *x_i_.* (2) is then updated to,

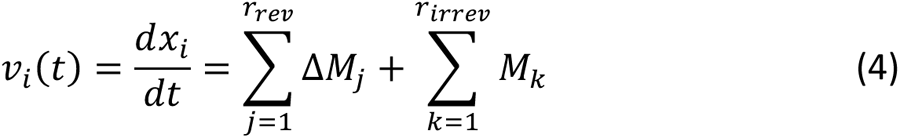

where the first sum is over reversible reactions (*r*_rev_) and the second sum is over irreversible reactions (*r_irrev_*).

We now define the set of reaction rate terms from (4) as:

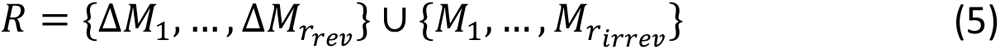

Since we only want to trace back the production of species *x_i_*, we can reduce *R* to its subset containing only the positive terms *R*_+_,

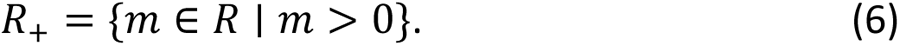

The most dominant monomial contributing to the production of *x_i_* is then given by,

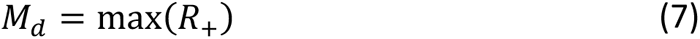

where *M*_d_ identifies the most dominant reaction in the production of *x*_i_. Next, we want to identify any additional reactions which contribute to the production of *x*_i_ with a similar magnitude as *M*_d_. We, therefore, apply the concept of dominancy as defined by Noel et al. (*31*), where monomials *M*_i_and *M*_j_ are said to be on par with each other within a level ρ > 0 if

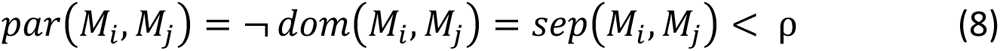

where *dom*(*M*_i_, *M*_j_) is a binary function of monomials for which monomial *M_j_* is said to dominate monomial *M_k_* at a level ρ > 0 if,

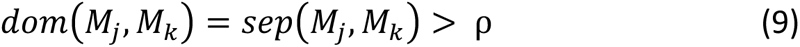

and *sep*(*M_j_*, *M*_k_) is the separation (i.e., Euclidean distance) in logarithmic space between the monomials,

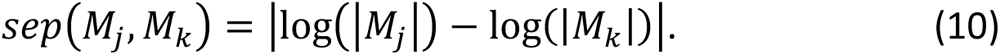

Using this application of dominancy (8), the set of dominant reactions terms, *D_xi_*, is given by,

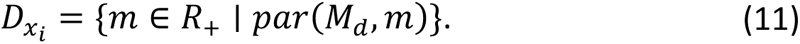

Thus, we construct a set of reaction terms containing the most dominant reaction term *M_d_* and all terms that are on par with *M_d_*. The level of separation ρ used to determine whether terms are on par is a user-defined quantity.

### 4. Obtaining modes of signal execution

Simulating biochemical models with different initial protein levels and/or kinetic parameters may result in different digital signatures. It is important to quantify the level of discrepancy between different sequences. Grouping sequences that have similar dominant pathways leads to the definition of modes of signal execution. First, it is necessary to find a suitable distance metric that accounts for the differences of interest when comparing sequences. Once we have a metric, we can obtain a distance matrix that can be used with clustering algorithms to identify the modes of signal execution.

#### a. Defining a suitable distance measure for clustering

Multiple distance measures exist that can be used the calculate the dissimilarity between two sequences. Each of the distances has different sensitivities to sequencing, timing, and duration of a dominant subnetwork. Since our goal is to identify groups of simulations that have the same dominant subnetwork, we chose the Longest Common Subsequence (LCS) distance (*42*). This metric is more sensitive to sequencing, i.e., the order of appearance of dominant paths in a signature. By focusing on the differences in the state distribution, it allows us to identify different modes of signal execution and novel protein targets that modulate biochemical signals within a network depending on the parameters of the model.

The LCS distance is defined as:

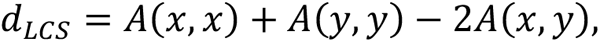

where *A*_s_(*x*, *y*) corresponds to the number of elements in one sequence that can be uniquely matched with elements occurring in the same order (not necessarily contiguous) in the other sequence.

We apply this metric to all pairs of digital signatures to obtain a dissimilarity matrix. Then, we use this matrix as an input of a clustering technique to obtain groups of digital signatures that have similar patterns in the sequence of dominant pathways. There are three clustering techniques implemented: Agglomerative (*44*), spectral (*66*), and HDBSCAN clustering (*67*). The choice of clustering technique is defined by the user.

### 5. Reduction of execution mode uncertainty

#### a. XGBOOST model of execution mode estimation

We used XGBOOST (*68*) to classify parameter vectors into different execution methods. We used 75%, and 25% of the parameter vectors as training and test sets, respectively. We used the following parameters to run the XGBOOST model: booster=’gbtree’, alpha=1, lambda=1, tree_method=’gpu_hist’, tree_method=’gpu_hist’, max_bin=16, objective=’multi:softmax’, num_class=12, eval_metric=’merror’, learning_rate=0.1, n_estimators=100.

To obtain the importance of individual parameters for the classification we used the ‘total_gain’ variable from the XGBOOST model.

#### b. Bayesian update of mode probabilities

After measurement of a kinetic parameter, mode probabilities may be updated using the following equation:

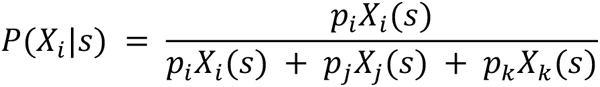

where *s* is the measured value, *X_i_* is the probability density function of the execution mode *i*, *X_j_* and *X_k_* are the probability density functions of the remaining two execution modes, and *P_i_*, *P_j_* and *p_k_* are their corresponding prior probabilities.

## Acknowledgements

The authors would like to thank Erin Shockley, Alexander Lubbock, Leonard Harris, Vito Quaranta, and Eric Deeds for insightful conversations and critical feedback on this work.

## Author contributions

**O.O.O.** developed the methods, performed the simulations and computations, and wrote the manuscript.

**M.O.** developed the methods, performed the simulations and computations, and wrote the manuscript.

**B.A.W.** contributed to and discussed the research topics and edited the manuscript.

**J.C.P.** contributed to and discussed the research topics and edited the manuscript.

**M.W.I.** contributed to and discussed the research topics and edited the manuscript.

**G.V.I.** contributed to and discussed the research topics and edited the manuscript.

**S.P.G.** contributed to and discussed the research topics and edited the manuscript.

**C.F.L.** conceived the ideas and concepts, developed the methods, and wrote the manuscript.

## Data and materials availability

All the code to reproduce the figures that contain model calibration, modes of signal execution, visualizations, and hypothesis exploration, is open source and can be found as Jupyter Notebooks in this GitHub repository: https://github.com/LoLab-VU/pydyno. These shareable and reusable notebooks contain all the source code and markup text that explains the rationale for each step in the analysis.

