## Supplemental information for "Emergent signal execution modes in biochemical reaction networks calibrated to experimental data"

Contents

1. Reducing execution mode uncertainty through parameter measurements
2. Modes of signal execution in a detailed apoptosis model with increased parameter uncertainty
3. Supplementary Tables S1-S3
4. Supplementary Figures S1-S11
5. Supplementary Videos S1-S11

### 1 Reducing execution mode uncertainty through parameter measurements

Given that the aEARM calibrated parameter vectors yield three execution modes (Figure 2C-D in the main text) with their respective probabilities, there is uncertainty about which execution mode is most representative of the cellular process. We then asked whether we could identify parameters that, if measured experimentally, would reduce the execution mode uncertainty. We hypothesize that identifying key parameters that inform execution mechanisms could guide experiments to improve our knowledge about network-driven signal processing. To measure the uncertainty of the execution modes, we used Shannon's entropy[3] formulated as $H = -\sum_{i=1}^{n} P(x_i) \log_2 P(x_i)$. As aEARM has three execution modes, the maximum entropy in the system is $\log_2 3 = 1.58$, which would signify that each execution mode has a 33% probability. Using the probabilities of the previously obtained execution modes (Figure 2C in the main text) and Shannon's formula, we calculated an entropy of 1.54 indicating a high uncertainty in the execution across all modes. To determine the most informative parameters that should be measured to reduce execution mode uncertainty, we used XGBOOST[2], a gradient-boosted Machine Learning technique that can classify parameter vectors into their corresponding execution modes. We used the calibrated parameter sets as training data where each kinetic parameter (kf, kr, kc, i.e., forward, reverse, and catalysis reaction constants, respectively) is a feature, and the mode of execution is our target variable.

Feature importance analysis from the XGBOOST analysis shows that the model parameters kf6 and kf7 contribute the most to training loss reduction during the classification task (Figure S6A). As illustrated in Figure S6C, parameters kf6 and kf7 correspond to the binding rate of MOMP* to inactivated MOMP, and MOMP* to eC, respectively. These two parameters are part of the reactions where the signal flux is bifurcated in the network, indicating that their values play an important role in the definition of the execution modes. To show the differences in parameter values for each execution mode we plotted the values of the kf6 and kf7 parameters. As shown in Figure S6B the execution modes have different distributions of the kf6 and kf7 parameters with some overlap. To show how measuring the kinetic parameters, the binding rate of MOMP* to eC, reduces the uncertainty in the mode of signal execution, we simulated 100 measurements of the kf7 parameter, and calculated the entropy with the updated execution modes probabilities. As shown in Figure S6D, measurements in the range of -8.1 and -7.15, where there is no overlap between the parameter distributions, reduce the entropy to less than 0.05 which indicates that we know the execution mode with certainty. For measurements larger than -7.15 but lower than -5.37 the overlap between distributions starts to increase and with it the uncertainty increases as well, as indicated by the entropy. Then, for measurements larger than -5.37 the entropy starts to decrease. This indicates that in general measuring kinetic parameters reduces the entropy and thus the uncertainty in the execution mode with the exception where the overlap between distribution is maximum and increases the entropy.

#### 2 Modes of signal execution in a detailed apoptosis model with increased parameter uncertainty

Based on our results with aEARM, we then asked how a larger model with higher parameter uncertainty would fare under the presented signal execution analysis. We shifted to a larger extrinsic apoptosis reaction model (EARMv2.0), which has been studied and characterized in previous work[1]. As illustrated in Figure S7A, EARMv2.0 is considerably larger than aEARM as the biochemical interactions are described with higher molecular resolution. In all, EARMv2.0 has 77 molecular species and 105 kinetic parameters. As described in Materials and Methods, we used PyDREAM to calibrate the model to published experimental data[4]. Although the calibration yielded parameter vectors that fit the experimental data indistinguishably well (Figure S8), we note that only 62 model parameters converged according to the Gelman-Rubin diagnostic (GR < 1.2) after two million iterations (see Table S3 and Figure S9). Distributions of 9 converged parameters are shown in Figure S7B. The remaining parameters exhibited GR values between 1.21 and 13.52 (Table S2 and Figure S10a, b, and c). From a Bayesian perspective, non-convergent parameters imply that the experimental data simply cannot constrain their values to a distribution and thus results in higher variability. As our analysis is focused on understanding execution modes in network-driven processes, a model with poorly identified parameters presents an opportunity to explore how signal execution could be best interpreted and understood in large model systems with high parameter uncertainty.

We followed the same procedure used in the main text to explore the execution modes in EARMv2.0 (See Matherials and Methods for details). Our analysis shows that calibrating EARMv2.0 to experimental data constrains the signal flow to eleven execution modes, that can be represented by 53 dominant subnetworks out of 2067 possible subnetworks. As shown in Figure S7C, the apoptosis execution signal could flow through any of these paths with varying degrees of probability, with Execution Mode 1 (EM1) exhibiting a probability of ~20% and the first four modes capturing ~50% of the signal probability, thus suggesting high path entropy as we have seen in previous work[5]. Supplementary videos show animations of signal flow for all execution modes in the context of EARMv2.0. If one needs to compare EMs observed for aEARM and EARMv2.0, we see that there exists an overlap between the underlying highly probable dominant pathways although they visually seem different. The main difference in the EMs is that complex interactions in EARMv2.0 are coarse-grained into a single node in aEARM and thus the EMs appear simplified. For instance, the MOMP node in aEARM represents the mitochondrial outer membrane pore formation which is explicitly implemented in EARMv2.0 by the Bid to Smac pathways. Thus, the EMs obtained from aEARM and the EMs obtained from EARMv2.0 are comparable and comprise functional commonality. While the EARM models represent the same mechanism, their resolution and hence the underlying system of differential equations are different. Therefore, the functional commonality in the observed EMs may imply that the EMs are intrinsic features of the underlying system and independent of the specific way in which the model is implemented.

Next, we tested whether each execution mode exhibits different responses to the same perturbation. We selected EM1 and EM2 for analysis as these modes exhibit the highest probability for signal execution. As illustrated in Figure S7D, the mBid interaction with Mcl1 is dominant in EM1. In contrast, the mBid interactions with both Mcl1 and Bcl2 are dominant in EM2, thus highlighting the importance of both antiapoptotic proteins to understand the signaling mechanisms during apoptosis execution when the cell is exposed to an apoptotic inducer.

We then performed two *in silico experiments* for EM1 and EM2: (i) a 50% knockdown (KD) of the antiapoptotic protein Mcl1 as well as (ii) a 50% knockdown of the antiapoptotic protein Bcl2. For the Mcl1 KD, we found that the EM1 median Time of Death (ToD) decreased from 10022.46 s (WT) to 8686.52 s (Figure S11A-upper panel). This is expected since mBid and Mcl1 interactions are dominant in this mode. By contrast the median ToD in EM2 decreased from 9943.65 s (WT) to 9335.85 s. This modest decrease in ToD can be attributed to the fact that although Mcl1 and mBid interactions are important in EM2, the dominance of Bcl2 compensates for the absence of Mcl1 and reduces the impact on ToD for the Mcl1 KD.

For the Bcl2 KD, we found that the median ToD in EM1 has a minor change from 10022.46 s to 10011.87 s, as expected because mBid activity is not significantly affected by Bcl2 in this mode. By contrast, in EM2, mBid activity is modulated by Mcl1 and Bcl2. Thus, a reduction in the initial protein levels of Bcl2 enables more mBid proteins to activate pro-apoptotic proteins and this leads to an increase in ToD to 9580 s (Figure S11A-upper panel). Taken together this data shows that distinct execution modes respond differently to the same perturbation and that their responses can be predicted based on the dominant reactions for a given execution mode.

To further emphasize the importance of transient dynamics on signal processing, we explored EM1 dynamic fingerprints and found that SMAC inhibition of XIAP occurs at later time points of the simulations (>8640 s). Therefore, we hypothesized that XIAP inhibition would be more effective earlier during signal execution. To test this, we added an XIAP inhibitor to EARMV2.0 at either 4000 s or 8000 s. As shown, when the inhibitor is added at the later time point, we observed a small reduction (Figure S11A lower panel) in the median ToD from 9943.65 s in the WT to 9380.44 s ($\Delta t$ = 563.21 s). In contrast, when the inhibitor is added at the earlier time point, when SMAC is not yet released from the mitochondria, the inhibitor binds to XIAP enabling C3 to cleave PARP and thereby reducing the median ToD to 6766.10 ($\Delta t$ = 3177.55 s).

As combination therapies have become important to combat drug resistance[6,7], especially in cancers, we explored whether our analysis provided information about potential targets for co-treatment. As we previously mentioned, Mcl1 and XIAP are dominant antiapoptotic proteins in EM1, thus we hypothesized that inhibition of both proteins would yield a shorter ToD compared to only inhibiting XIAP. To test this, we added two drugs that independently inhibit XIAP and Mcl1 and obtained a ToD of 5951.11 s representing a 12% reduction in the ToD compared to XIAP inhibition only (Figure S11A lower panel). Finally, to guide experiments that would identify the most likely execution mode out of the 11 execution modes obtained, we developed an XGBOOST model of execution mode estimation and performed feature importance analysis. As shown in Figure S11B, we found that the parameters controlling the kinetics of mBid binding to BcxL, and XIAP binding to C3 yield the most information about execution modes in EARMv2.0. Taken together, these results suggest that the analysis of signaling dynamics from uncertain parameters helps us identify dominant reactions that control signal flow in a network during signal processing and how these networks are more sensitive to the perturbations of those reactions.

### 3 Supplementary Tables

*Table S1. Gelman-Rubin (GR) diagnostic values for all calibrated parameters in the aEARM model. Parameters with GR values lower than 1.2 are considered to be converged*

| Parameter | GR |
|---|---|
| kf0 | 1.08514627 |
| kr0 | 1.00220123 |
| kc0 | 1.03226655 |
| kf1 | 1.0526068 |
| kr1 | 1.00504595 |
| kc1 | 1.02257591 |
| kf2 | 1.0130557 |
| kr2 | 1.00305479 |
| kc2 | 1.00393523 |
| kf3 | 1.00236732 |
| kr3 | 1.00518277 |
| kc3 | 1.00208664 |
| kf4 | 1.03636438 |
| kr4 | 1.00973982 |
| kc4 | 1.07074804 |
| kf5 | 1.06039857 |
| kr5 | 1.11159688 |
| kc5 | 1.06253039 |
| kf6 | 1.16079145 |
| kr6 | 1.01920037 |
| kc6 | 1.12806401 |
| kf7 | 1.1978977 |
| kr7 | 1.015523 |
| kc7 | 1.04973795 |
| kf8 | 1.09112645 |
| kr8 | 1.00130727 |
| kc8 | 1.05073858 |
| kc9 | 1.00629721 |

*Table S2. Silhouette score for different numbers of clusters for aEARM.*

| Number of clusters | Silhouette Score |
|---|---|
| 2 | 0.696260 |
| **3** | **0.935141** |
| 4 | 0.886408 |
| 5 | 0.872150 |
| 6 | 0.862335 |
| 7 | 0.828153 |
| 8 | 0.827130 |
| 9 | 0.814062 |
| 10 | 0.810126 |

*Table S3. Gelman-Rubin (GR) diagnostic values for all calibrated parameters in the EARMv2.0 model. Parameters with GR values lower than 1.2 are considered to be converged*

| Parameter | GR |
|---|---|
| bind_L_R_to_LR_kf | 1.30555024 |
| bind_L_R_to_LR_kr | 1.19366713 |
| convert_LR_to_DISC_kc | 1.59500279 |
| bind_DISC_C8pro_to_DISCC8pro_kf | 1.62566521 |
| bind_DISC_C8pro_to_DISCC8pro_kr | 1.01651954 |
| catalyze_DISCC8pro_to_DISC_C8A_kc | 1.00638202 |
| bind_C8A_BidU_to_C8ABidU_kf | 1.57868642 |
| bind_C8A_BidU_to_C8ABidU_kr | 1.03076058 |
| catalyze_C8ABidU_to_C8A_BidT_kc | 6.63072247 |
| bind_DISC_flip_kf | 1.02016009 |
| bind_DISC_flip_kr | 1.02119955 |
| bind_BAR_C8A_kf | 2.14919922 |
| bind_BAR_C8A_kr | 1.00174068 |
| equilibrate_SmacC_to_SmacA_kf | 1.01189403 |
| equilibrate_SmacC_to_SmacA_kr | 1.03203665 |
| equilibrate_CytoCC_to_CytoCA_kf | 1.1986676 |
| equilibrate_CytoCC_to_CytoCA_kr | 1.12600228 |
| bind_CytoCA_ApafI_to_CytoCAApafI_kf | 1.22739716 |
| bind_CytoCA_ApafI_to_CytoCAApafI_kr | 1.02685524 |
| catalyze_CytoCAApafI_to_CytoCA_ApafA_kc | 1.03909011 |
| convert_ApafA_C9_to_Apop_kf | 2.04271115 |
| convert_ApafA_C9_to_Apop_kr | 1.45004061 |
| bind_Apop_C3pro_to_ApopC3pro_kf | 3.88094106 |
| bind_Apop_C3pro_to_ApopC3pro_kr | 1.46376778 |
| catalyze_ApopC3pro_to_Apop_C3A_kc | 2.12805512 |
| bind_Apop_XIAP_kf | 1.63051009 |
| bind_Apop_XIAP_kr | 1.26680774 |
| bind_SmacA_XIAP_kf | 1.05473289 |
| bind_SmacA_XIAP_kr | 1.0034777 |
| bind_C8A_C3pro_to_C8AC3pro_kf | 3.43426491 |
| bind_C8A_C3pro_to_C8AC3pro_kr | 1.01372768 |
| catalyze_C8AC3pro_to_C8A_C3A_kc | 1.87635914 |
| bind_XIAP_C3A_to_XIAPC3A_kf | 1.67972634 |
| bind_XIAP_C3A_to_XIAPC3A_kr | 1.15656166 |

| | |
|---|---|
| catalyze_XIAPC3A_to_XIAP_C3ub_kc | 1.09385208 |
| bind_C3A_PARPU_to_C3APARPU_kf | 1.25018223 |
| bind_C3A_PARPU_to_C3APARPU_kr | 1.04884998 |
| catalyze_C3APARPU_to_C3A_PARPC_kc | 1.12017976 |
| bind_C3A_C6pro_to_C3AC6pro_kf | 13.5217835 |
| bind_C3A_C6pro_to_C3AC6pro_kr | 1.03639819 |
| catalyze_C3AC6pro_to_C3A_C6A_kc | 1.03041378 |
| bind_C6A_C8pro_to_C6AC8pro_kf | 4.69722606 |
| bind_C6A_C8pro_to_C6AC8pro_kr | 1.07790728 |
| catalyze_C6AC8pro_to_C6A_C8A_kc | 1.120102 |
| equilibrate_BidT_to_BidM_kf | 2.61600196 |
| equilibrate_BidT_to_BidM_kr | 3.09256962 |
| bind_BidM_BaxC_to_BidMBaxC_kf | 3.70912317 |
| bind_BidM_BaxC_to_BidMBaxC_kr | 1.02294992 |
| catalyze_BidMBaxC_to_BidM_BaxM_kc | 1.07300834 |
| bind_BidM_BaxM_to_BidMBaxM_kf | 1.23579073 |
| bind_BidM_BaxM_to_BidMBaxM_kr | 1.02038414 |
| catalyze_BidMBaxM_to_BidM_BaxA_kc | 1.08771274 |
| bind_BidM_BakM_to_BidMBakM_kf | 1.05678249 |
| bind_BidM_BakM_to_BidMBakM_kr | 1.18477726 |
| catalyze_BidMBakM_to_BidM_BakA_kc | 1.29135272 |
| bind_BaxA_BaxM_to_BaxABaxM_kf | 1.44060819 |
| bind_BaxA_BaxM_to_BaxABaxM_kr | 1.00404728 |
| catalyze_BaxABaxM_to_BaxA_BaxA_kc | 1.02261922 |
| bind_BakA_BakM_to_BakABakM_kf | 1.79854806 |
| bind_BakA_BakM_to_BakABakM_kr | 1.24688942 |
| catalyze_BakABakM_to_BakA_BakA_kc | 1.04152718 |
| bind_BidM_Bcl2M_kf | 1.32171588 |
| bind_BidM_Bcl2M_kr | 1.25176661 |
| bind_BidM_BclxLM_kf | 1.42163672 |
| bind_BidM_BclxLM_kr | 1.25451231 |
| bind_BidM_Mcl1M_kf | 2.52365114 |
| bind_BidM_Mcl1M_kr | 1.53131431 |
| bind_BaxA_Bcl2_kf | 1.03114713 |
| bind_BaxA_Bcl2_kr | 1.01410852 |
| bind_BaxA_BclxLM_kf | 1.48716584 |
| bind_BaxA_BclxLM_kr | 1.8534663 |
| bind_BakA_BclxLM_kf | 1.09290472 |

| | |
|---|---|
| bind_BakA_BclxLM_kr | 1.28694699 |
| bind_BakA_Mcl1M_kf | 1.03932055 |
| bind_BakA_Mcl1M_kr | 1.01668657 |
| bind_BadM_Bcl2_kf | 1.00495924 |
| bind_BadM_Bcl2_kr | 1.01024336 |
| bind_BadM_BclxLM_kf | 1.01820425 |
| bind_BadM_BclxLM_kr | 1.01971635 |
| bind_NoxaM_Mcl1M_kf | 1.01150003 |
| bind_NoxaM_Mcl1M_kr | 1.02564465 |
| assemble_pore_sequential_Bax_2_kf | 1.99372138 |
| assemble_pore_sequential_Bax_2_kr | 1.0432823 |
| assemble_pore_sequential_Bax_3_kf | 1.0519527 |
| assemble_pore_sequential_Bax_3_kr | 1.00962278 |
| assemble_pore_sequential_Bax_4_kf | 1.013291 |
| assemble_pore_sequential_Bax_4_kr | 1.00694199 |
| assemble_pore_sequential_Bak_2_kf | 1.24674091 |
| assemble_pore_sequential_Bak_2_kr | 1.05713598 |
| assemble_pore_sequential_Bak_3_kf | 1.01164877 |
| assemble_pore_sequential_Bak_3_kr | 1.03205593 |
| assemble_pore_sequential_Bak_4_kf | 1.0717765 |
| assemble_pore_sequential_Bak_4_kr | 1.00338011 |
| pore_transport_complex_BaxA_4_CytoCM_kf | 1.36126051 |
| pore_transport_complex_BaxA_4_CytoCM_kr | 1.0860153 |
| pore_transport_dissociate_BaxA_4_CytoCC_kc | 1.19093017 |
| pore_transport_complex_BaxA_4_SmacM_kf | 1.21029872 |
| pore_transport_complex_BaxA_4_SmacM_kr | 1.02142634 |
| pore_transport_dissociate_BaxA_4_SmacC_kc | 1.08568557 |
| pore_transport_complex_BakA_4_CytoCM_kf | 1.1615882 |
| pore_transport_complex_BakA_4_CytoCM_kr | 1.04931225 |
| pore_transport_dissociate_BakA_4_CytoCC_kc | 1.22633259 |
| pore_transport_complex_BakA_4_SmacM_kf | 1.33965964 |
| pore_transport_complex_BakA_4_SmacM_kr | 1.02048459 |
| pore_transport_dissociate_BakA_4_SmacC_kc | 1.01618226 |

#### 5 Supplementary Figures

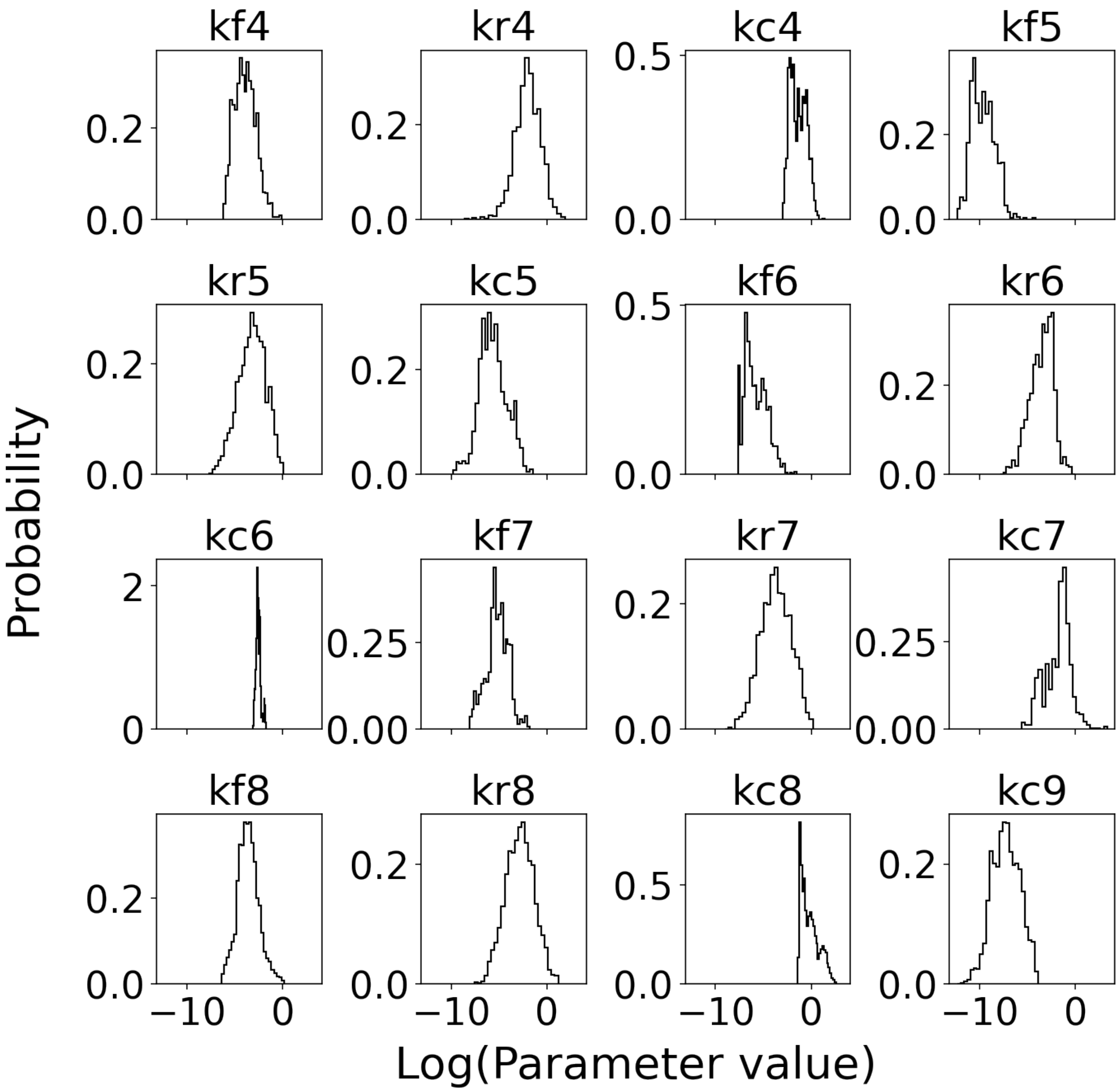


*Figure S1. Marginal probability distributions of aEARM individual kinetic parameters. Distributions that were recovered from the PyDREAM run by integrating out all other dimensions.*

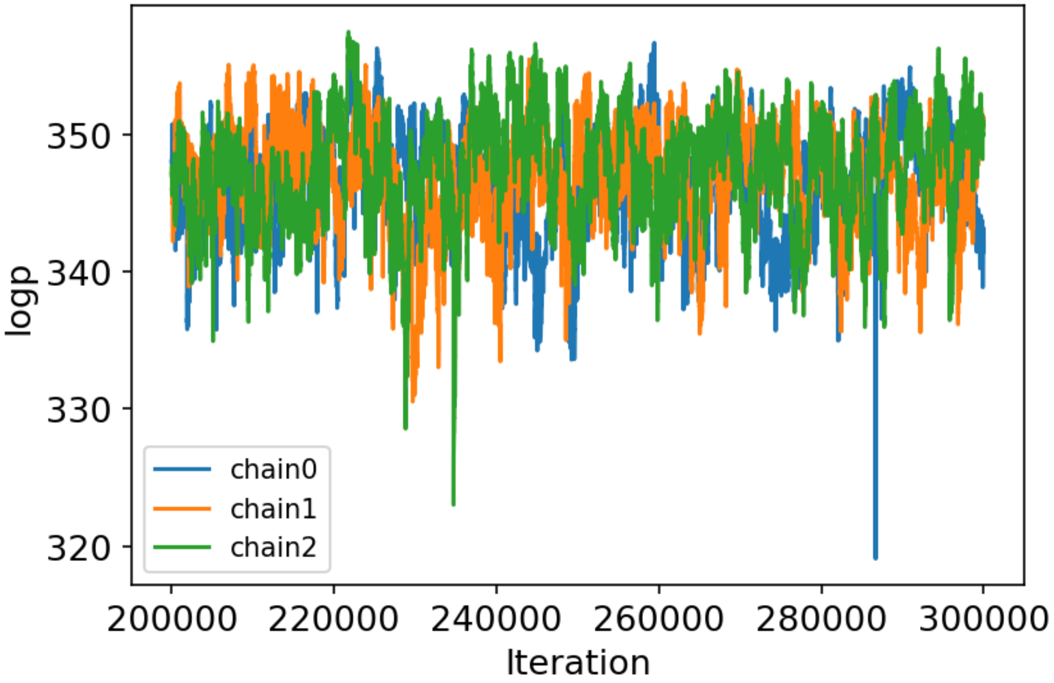


*Figure S2. Negative log-posterior values of the posterior distribution of aEARM sampled by 3 chains using PyDREAM after a burn-in of 200,000 iterations.*

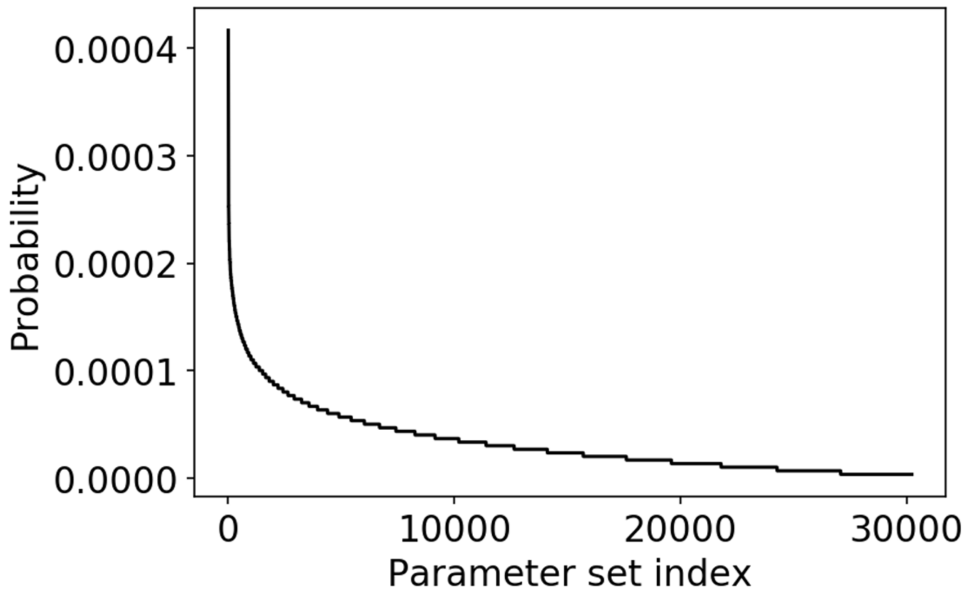


*Figure S3. Probability of each of the unique parameter vectors sampled after burn-in in the PyDREAM calibration. To obtain the probability of each parameter set the number of visits to a specific parameter vector was normalized by the total number of visits.*

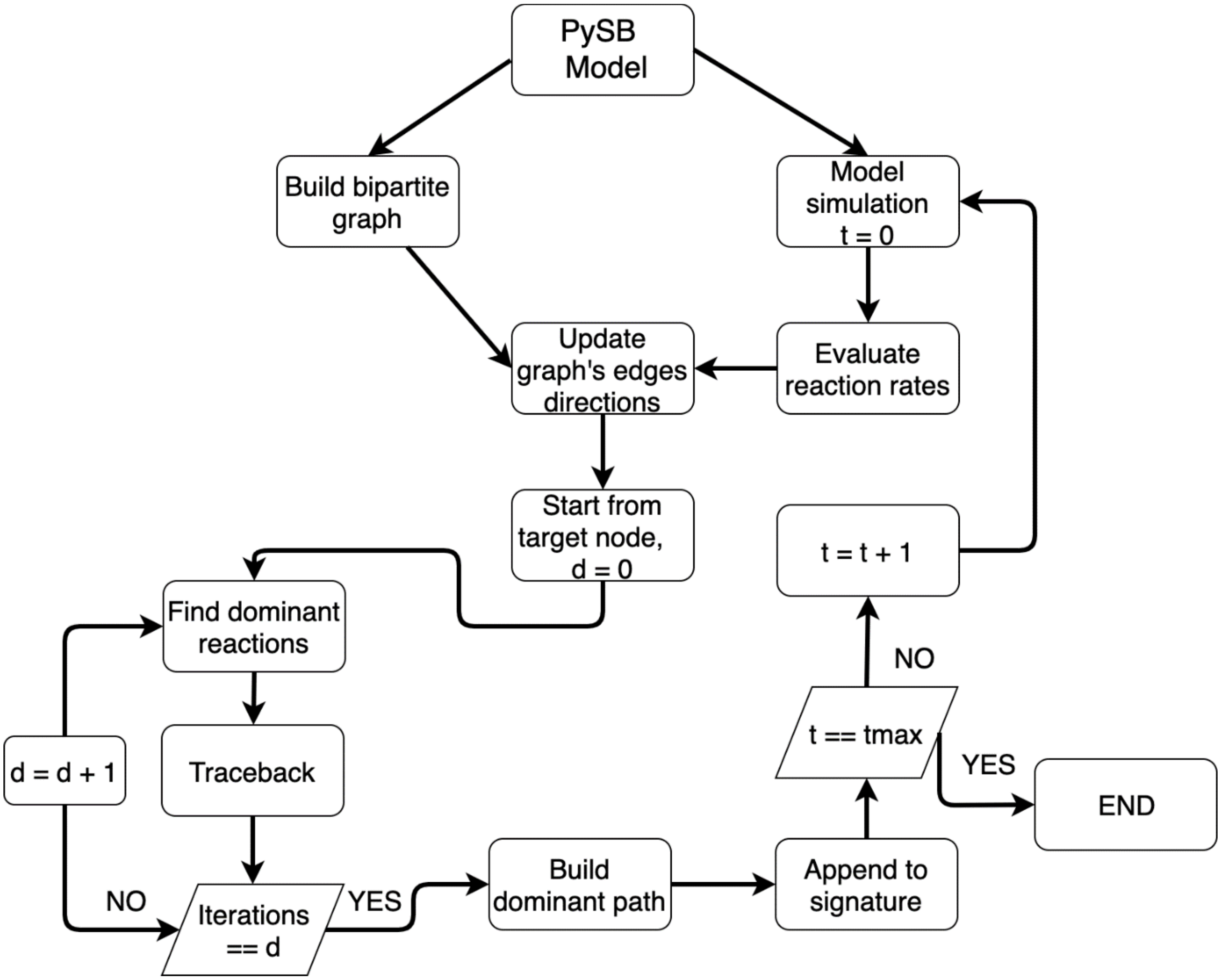


*Figure S4. Algorithm to obtain dynamic fingerprints from a simulation. Briefly, the algorithm consists in building a bipartite graph from the model, and then simulating the model with a specific parameter vector. Next, the algorithm obtains the dominant reactions from a network and builds a subnetwork. This procedure is repeated for each simulated time point.*

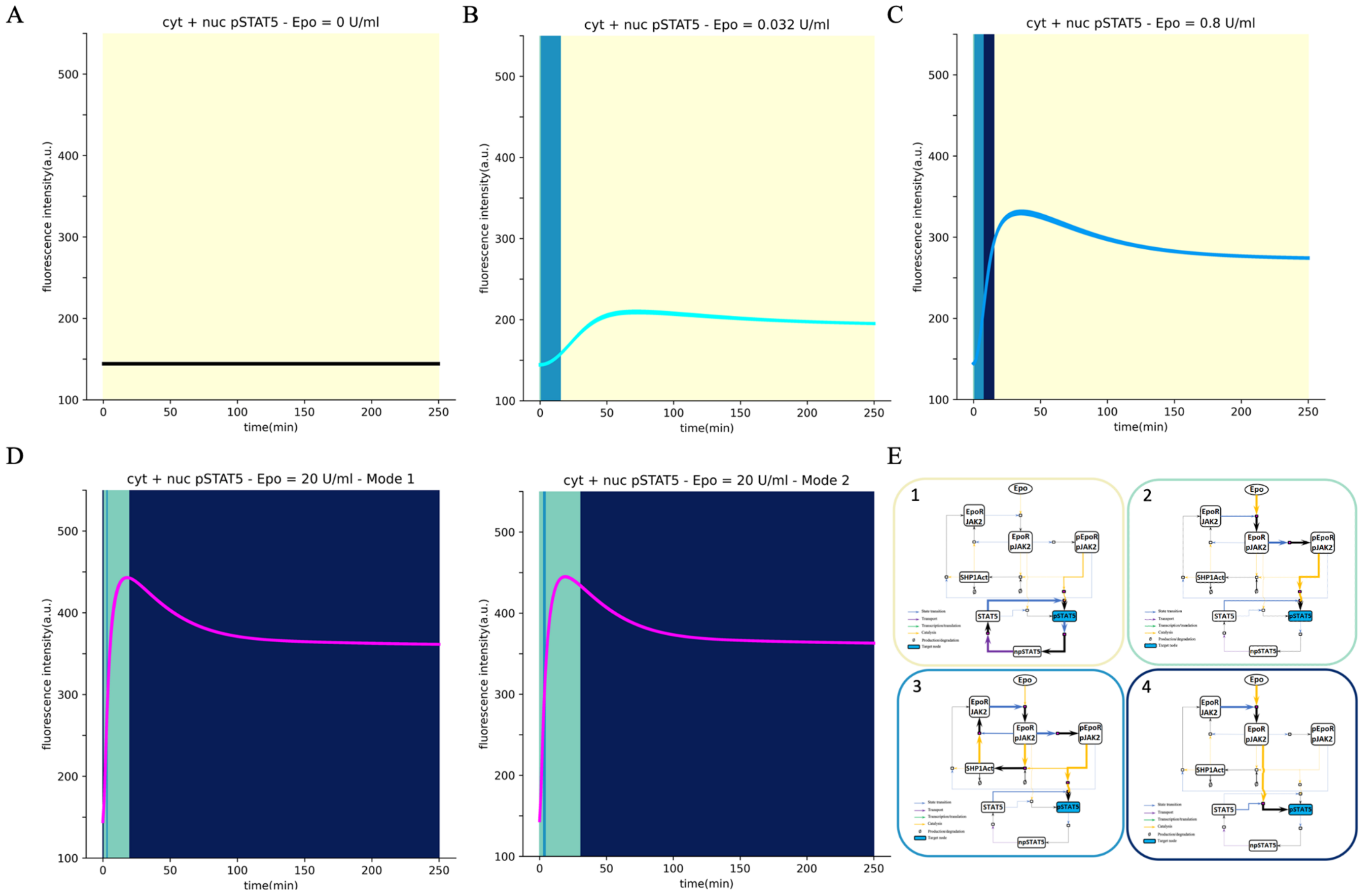


*Figure S5. Dominant pathways over time versus the pSTAT5 abundance with respect to increasing Epo doses. A, B, C, and D) Dominant pathways versus pSTAT5 trajectories when Epo = 0 U/ml, 0.032 U/ml, 0.8 U/ml, and 20 U/ml, respectively. E) Dominant pathways over time framed with the corresponding colors in A-D.*

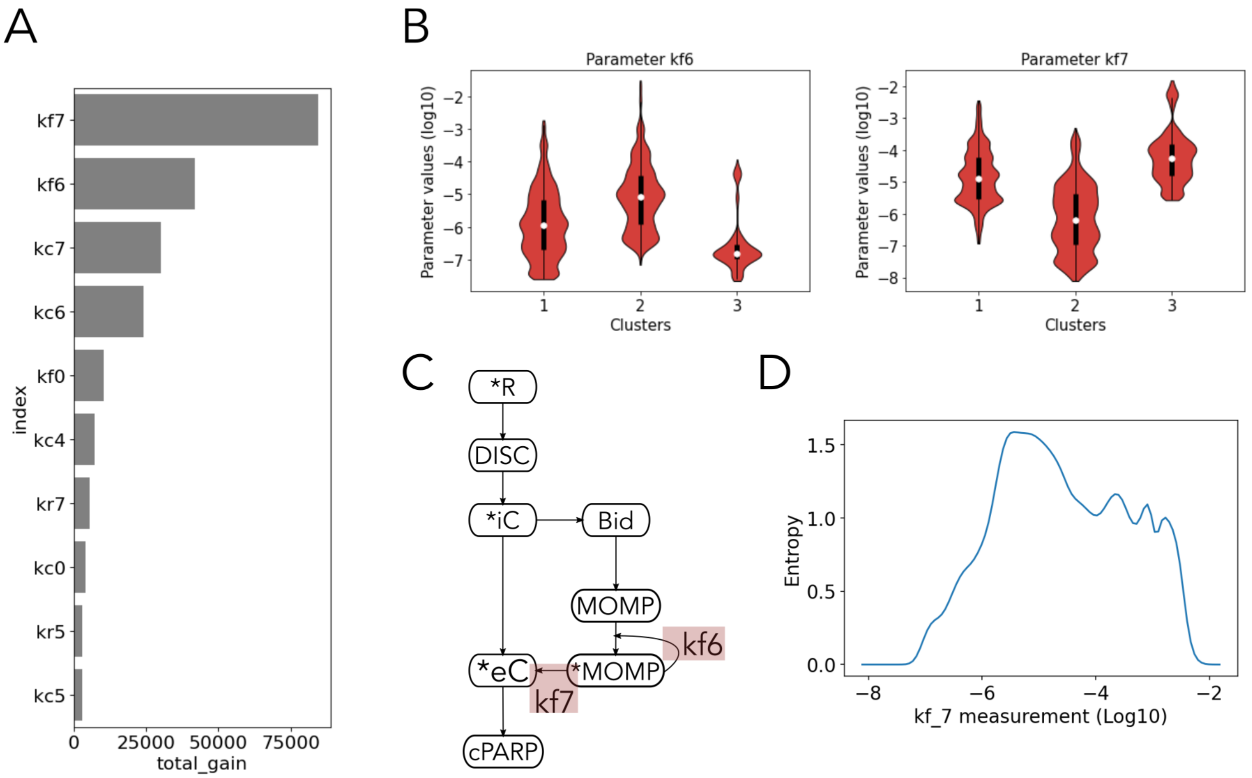


*Figure S6. Parameter measurements reduce aEARM execution mode uncertainty. A) List of the 10 parameters that contribute the most to model prediction. Parameters with higher total gains, compared to another parameter, provide larger improvements to accuracy in model prediction. B) Parameter values of kf6 and kf7 grouped by the execution mode they belong to. A Gaussian kernel was used to estimate the density probability of parameter values in each execution mode. C) Schematic representation of the aEARM network. Kinetic parameters kf6 and kf7 and their corresponding reactions are highlighted in the network. D) Changes in the execution modes entropy after simulated measurements of kf7.*

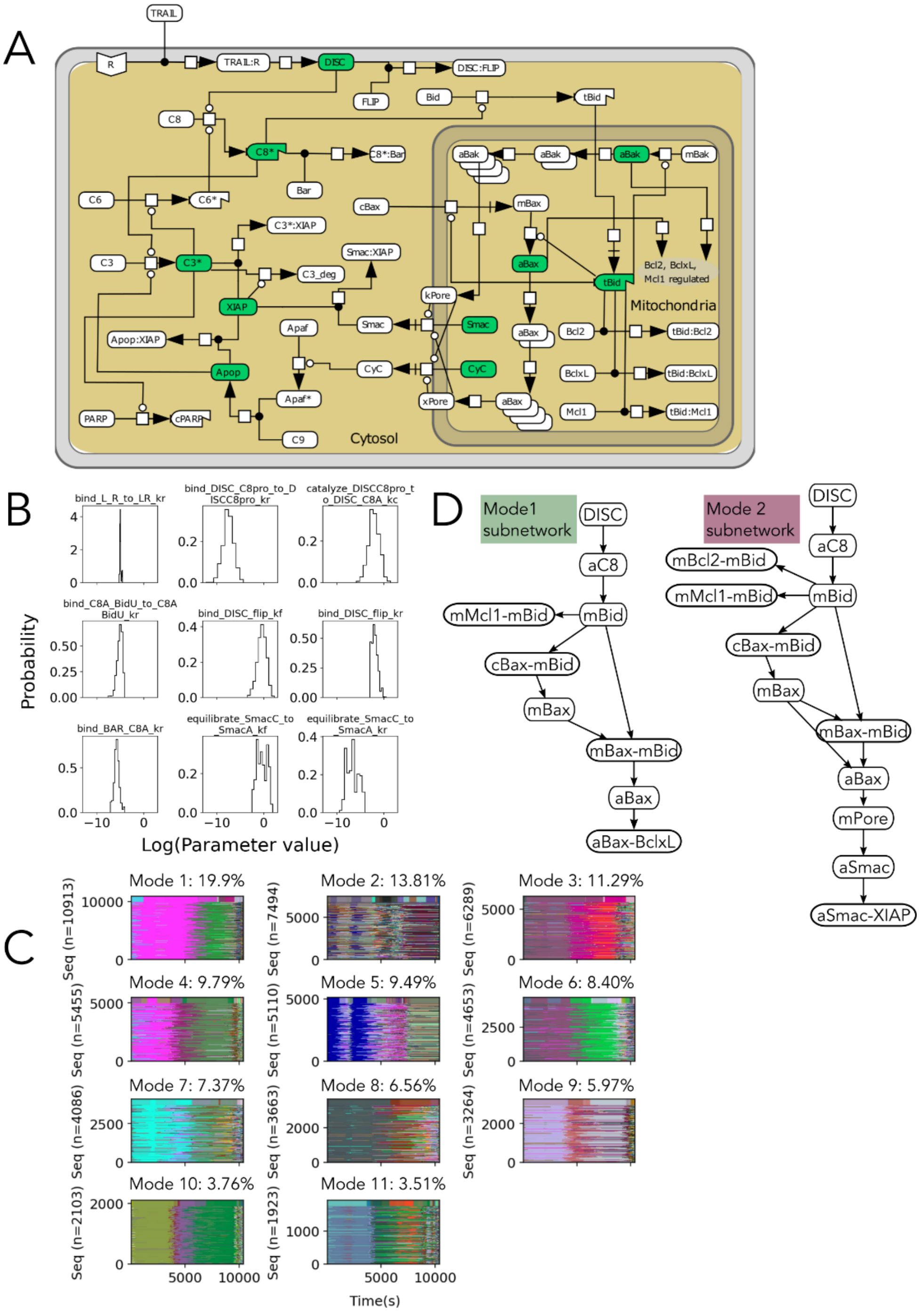


*Figure S7. Modes of signal execution in the full Extrinsic Apoptosis Reaction Network. A) Network of the interactions between the proteins in the apoptosis pathway. Proteins highlighted in green are nodes where the signal flux can be divided. The convention of Kitano[8] was followed. B) Marginal probability distributions of 9 individual kinetic parameters converged by the Gelman-Rubin diagnostic. C) Dynamic fingerprints organized by the execution modes they belong to. Each cluster plot is composed of horizontal lines that correspond to dynamic fingerprints, i.e. sequences of dominant subnetworks, and each subnetwork is assigned a different color. Execution modes are sorted from highest to lowest probability. D) Signal execution in Mode 1 (left) and Mode 2 (right) as defined by the most common subnetwork in each mode at t=7000s.*

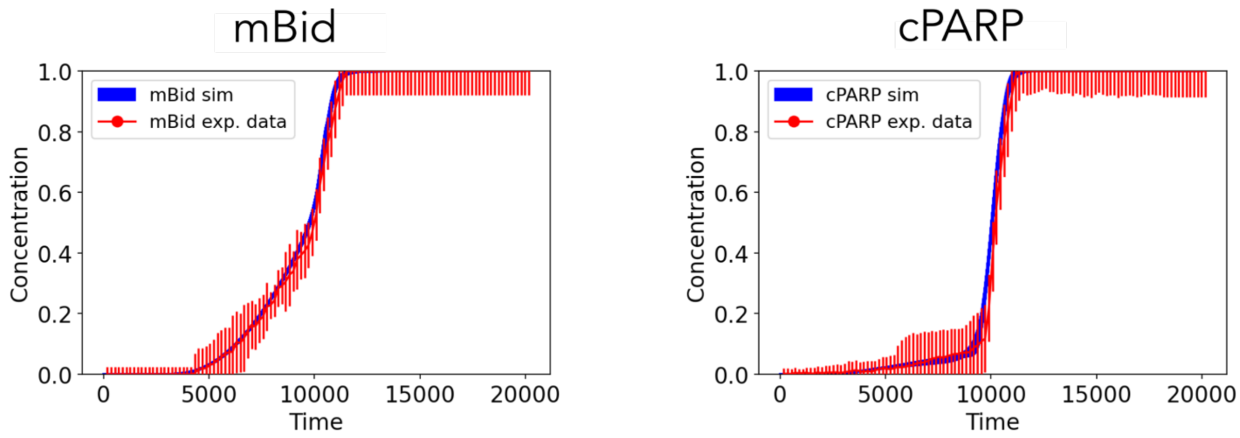


*Figure S8. Fit of EARMv2.0 to experimental data. Simulated trajectories of truncated Bid and Cleaved PARP calibrated to reproduce the experimental data. Red dots and bars indicate the mean and standard deviation of the experimental data and blue lines correspond to the simulated trajectories.*

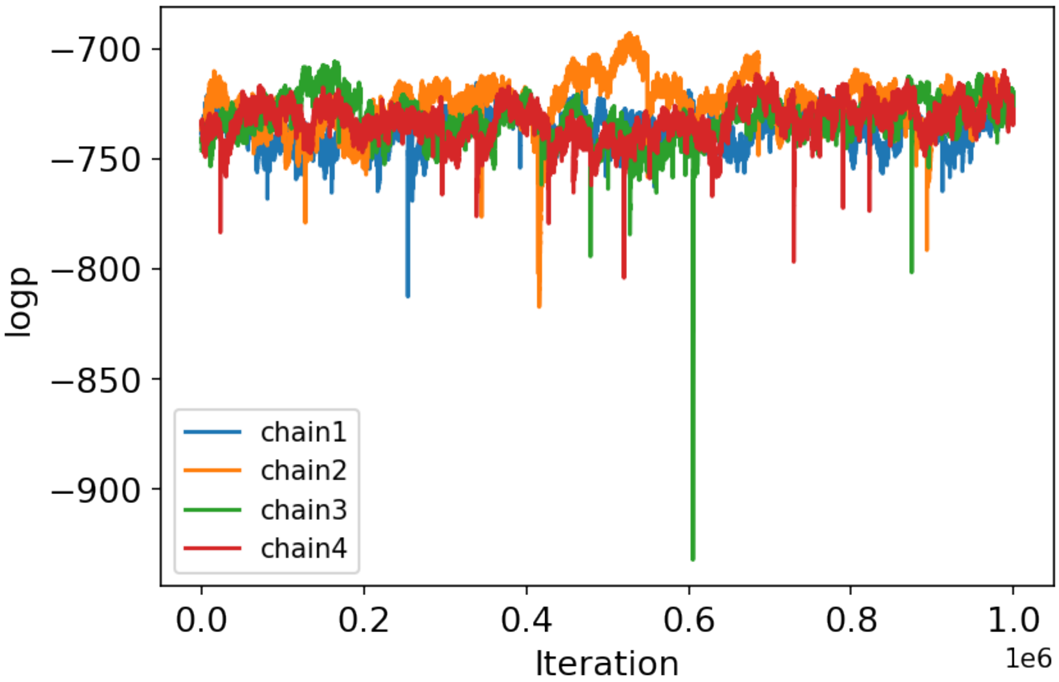


*Figure S9. Negative log-posterior values of the posterior distribution of EARMv2.0 sampled by 4 chains using PyDREAM after a burn-in of 1 million iterations.*

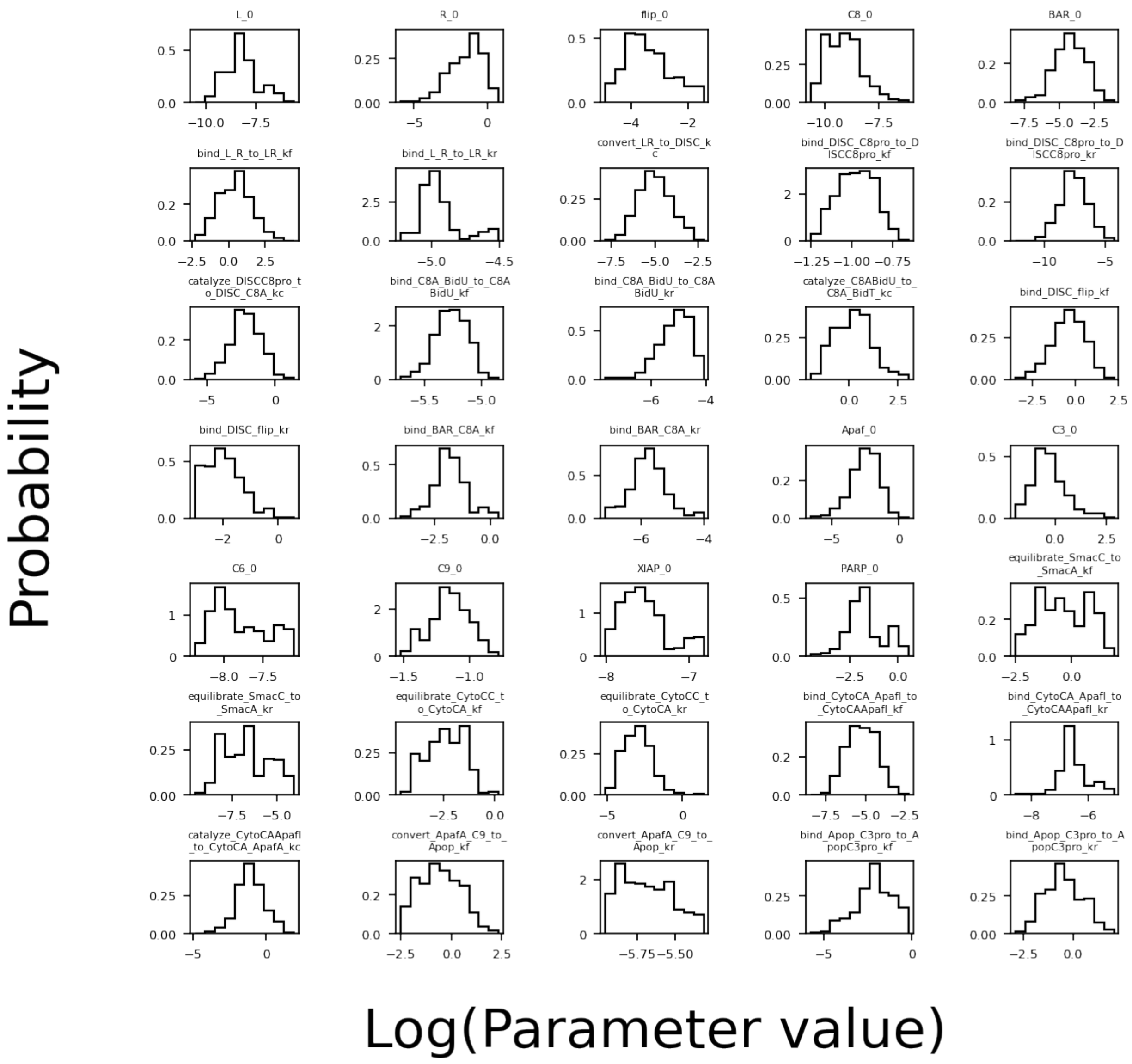


*Figure S10a. Marginal probability distributions of EARMv2.0 individual kinetic parameters. Distributions that were recovered from the PyDREAM run by integrating out all other dimensions.*

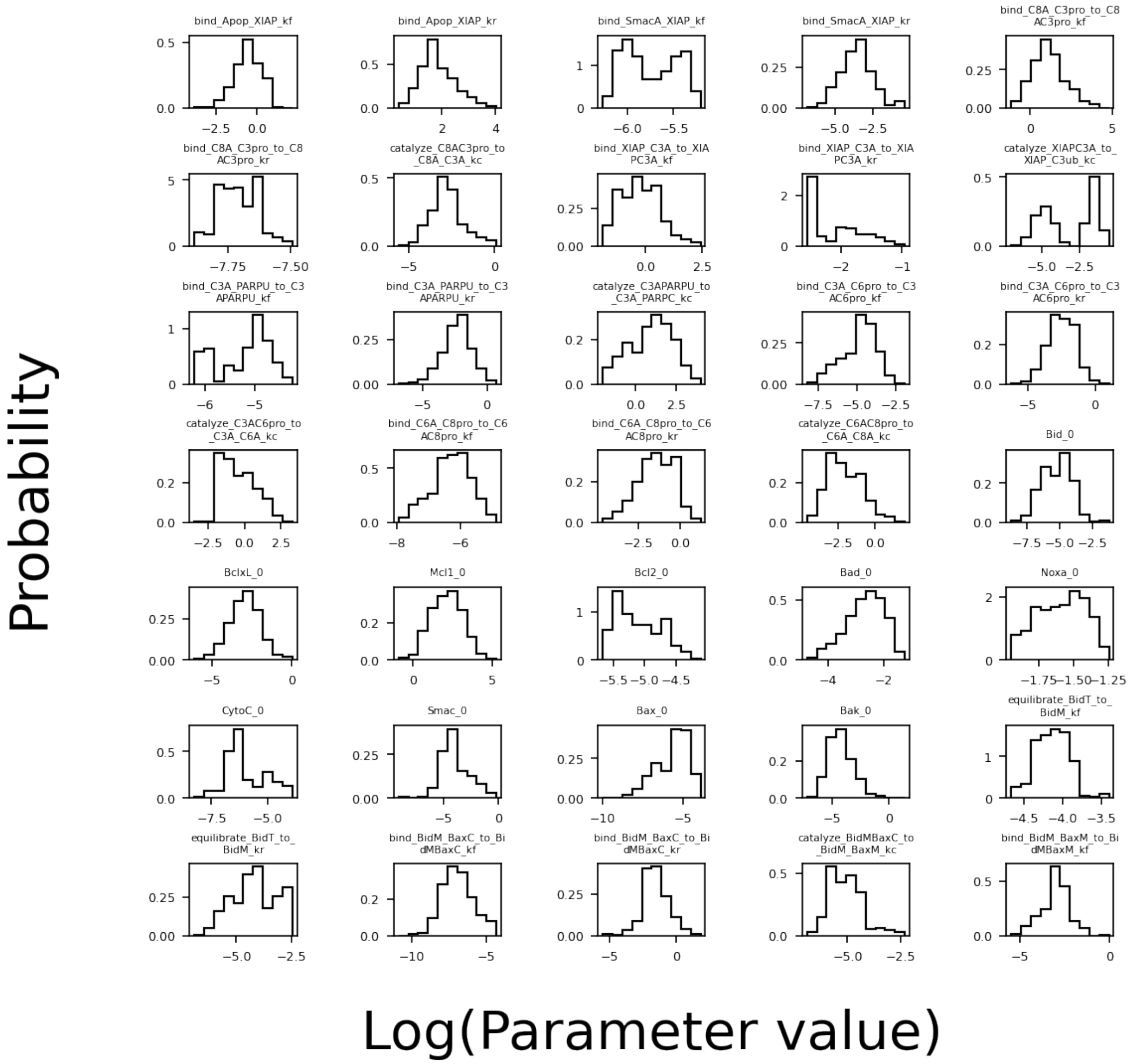


*Figure S10b. Marginal probability distributions of EARMv2.0 individual kinetic parameters. Distributions that were recovered from the PyDREAM run by integrating out all other dimensions.*

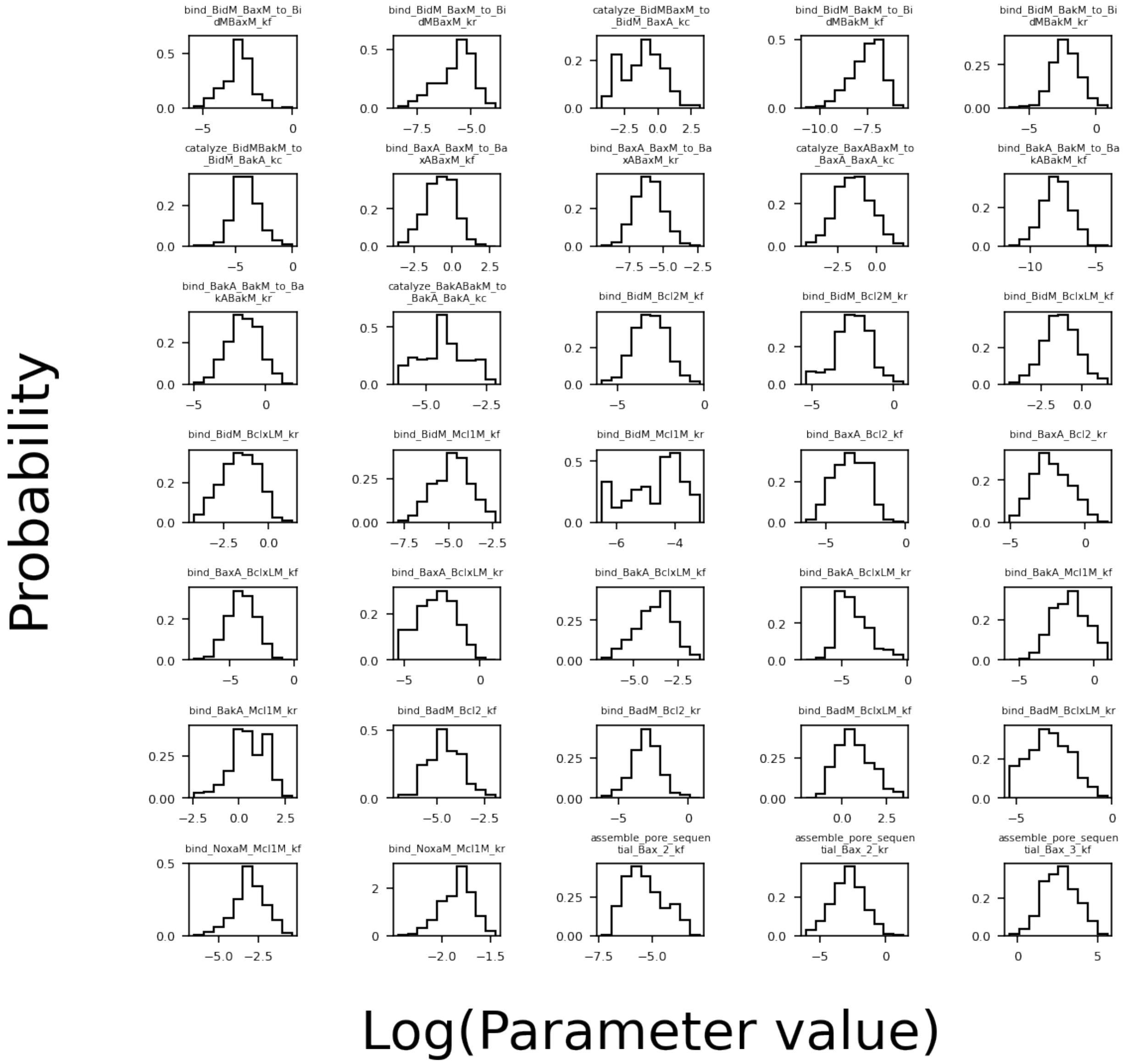


*Figure S10c. Marginal probability distributions of EARMv2.0 individual kinetic parameters. Distributions that were recovered from the PyDREAM run by integrating out all other dimensions.*

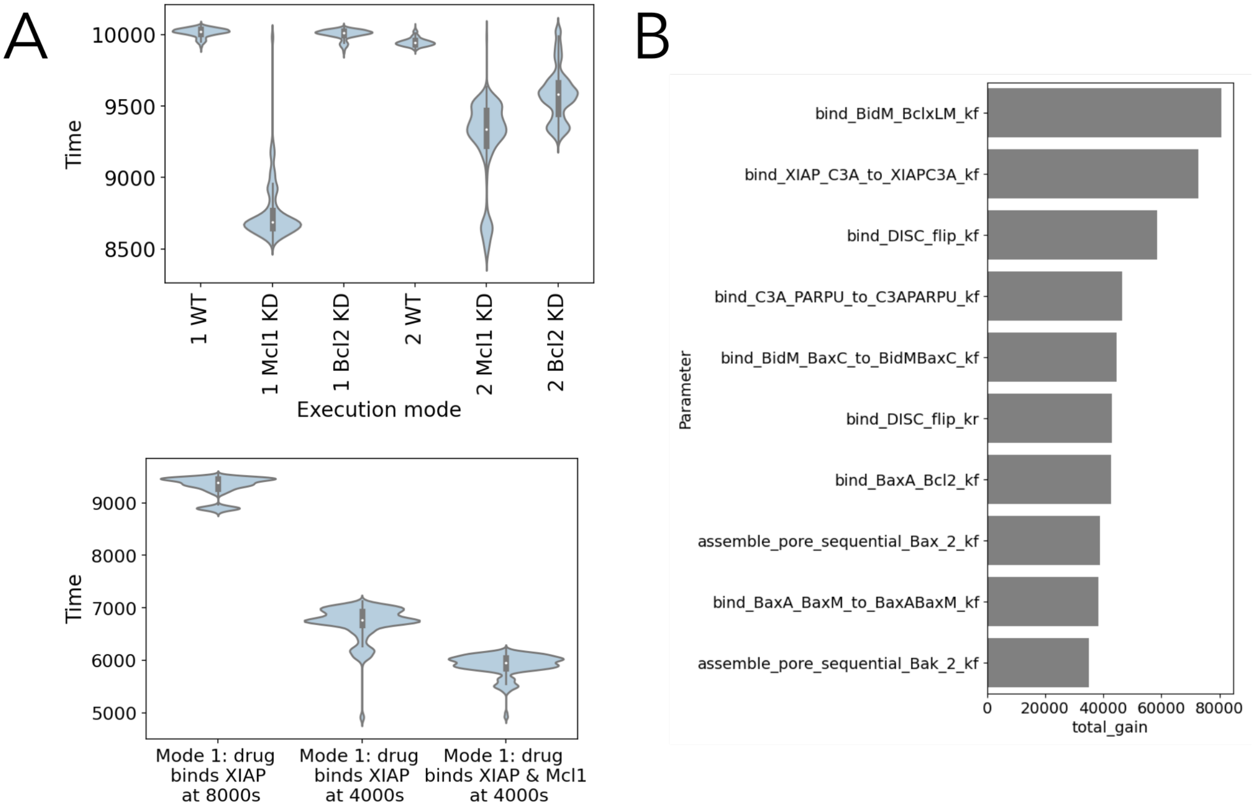


*Figure S11. In silico experiments to probe the different execution modes in EARMv2.0 A) Upper panel: Time to death distributions in the execution modes 1 and 2 in the "wild type" condition, after a 50% Mcl1 knockdown, and after a 50% knockdown of Bcl2. The boxplot inside the distributions shows the median, first quartile, and third quartile of the datasets. Execution modes 1 and 2 show substantial differences in their response to the knockdowns. Lower panel: Time to death distributions in the execution mode 1 after adding a drug that binds XIAP at t=8000 s, and at t=4000, and a drug that binds XIAP and Mcl1 at 4000 s. B) List of the 10 parameters in EARMv2.0 that contribute the most to the model prediction.*

### 6 Supplementary Videos:

Video S1-S11: Dynamic visualization of apoptosis execution for the 11 execution modes found in EARMv2.0 obtained using PyViPR[9]. Nodes represent molecular species, and pie charts inside the nodes depict relative species concentration over time. Edges represent the interaction between species, and shades of red show the intensity of the reactions.
